# Competition for fluctuating resources reproduces statistics of species abundance over time across wide-ranging microbiotas

**DOI:** 10.1101/2021.05.13.444061

**Authors:** Po-Yi Ho, Benjamin Good, Kerwyn Casey Huang

## Abstract

Across diverse microbiotas, species abundances vary in time with distinctive statistical behaviors that appear to generalize across hosts, but the origins and implications of these patterns remain unclear. Here, we show that many of these patterns can be quantitatively recapitulated by a simple class of resource-competition models, in which the metabolic capabilities of different species are randomly drawn from a common statistical ensemble. Our coarse-grained model parametrizes the intrinsic consumer-resource properties of a community using a small number of macroscopic parameters, including the total number of resources, typical resource fluctuations over time, and the average overlap in resource-consumption profiles across species. We elucidate how variation in these parameters affects various time series statistics, enabling macroscopic parameter estimation and comparison across wide-ranging microbiotas, including the human gut, saliva, and vagina, as well as mouse gut and rice. The successful recapitulation of time series statistics across microbiotas suggests that resource competition generally acts as a dominant driver of community dynamics. Our work unifies numerous time series patterns under one model, clarifies their origins, and provides a framework to infer macroscopic parameters of resource competition from longitudinal studies of microbial communities.

## Introduction

Microbial communities are ubiquitous across our planet, and strongly affect host and environmental health^1,2^. Predictive models for microbial community dynamics would accelerate efforts to engineer microbial communities for societal benefits. A well-studied class is consumer-resource (CR) models, wherein species growth is determined by the consumption of environmental resources^3^. CR models capture a core set of interactions among members of a community based on their competition for nutrients, and have demonstrated the capacity to recapitulate important properties of microbial communities such as diversity and stability^4,5^, despite neglecting potential interaction mechanisms beyond resource competition. Model parameters such as resource consumption rates are beginning to be uncovered in the context of *in vitro* experiments^6,7^. However, a systematic connection between CR models and complex *in vivo* systems is lacking.

A common strategy for interrogating *in vivo* microbiotas is longitudinal sampling followed by 16S amplicon or metagenomic sequencing, thereby generating a relative abundance time series. Analyses of longitudinal data have shown that species abundances fluctuate around stable, host-specific values in healthy humans^8-10^. Recently, it was discovered that such time series exhibit distinctive statistical signatures, sometimes referred to as macroecological dynamics^11-14^. For example, in human and mouse gut microbiotas, the variance versus mean abundance over time scales across species as a power law, recapitulating the empirical Taylor’s law observed in many ecosystems^15^. Taylor’s law and other statistical behaviors encode properties of the community and its environment; for example, deviations from Taylor’s law can highlight species that are transient invaders^11^. More broadly, the relative contributions of intrinsic versus environmental processes can be distinguished by modeling time series using autoregressive models whose output values depend linearly on values at previous times and external noise^16^. Abundance time series can be correlated to environmental metadata such as diet to generate hypotheses about how environmental perturbations affect community composition^10^. Time series analyses can also identify transitions between distinct ecological states^17^. Some statistical behaviors can be recapitulated by phenomenological models, such as a non-interacting, constrained random walk in abundances^12^ or a generalized Lotka-Volterra model with colored noise^13^. Similarly, ecological models describing the birth, immigration, and death of species have been used to analyze the distributions of abundance changes and abundance-prevalence relationships^18^. Nonetheless, the origins of and relationships among many of the statistical behaviors exhibited by host-associated microbiotas remain unclear.

Here we show that many of the observed statistical behaviors can be quantitatively recapitulated by a simple resource competition model, in which species abundance fluctuations arise solely from external fluctuations in resource levels. The resource consumption network of such a community will typically depend on thousands of underlying parameters. We sought to overcome this combinatorial complexity by adopting a coarse-grained approach, in which resources describe effective groupings of metabolites or niches, and model parameters are randomly drawn from a common statistical ensemble. We show that this simple model generates statistics that quantitatively match those observed in experimental time series across wide-ranging microbiotas, allowing us to infer the macroscopic, ensemble-level parameters of resource competition that can recapitulate their observed dynamics. Our work provides a systematic connection between complex microbiotas and the coarse-grained resource competition underlying their dynamics, with broad importance for engineering communities relevant to human health and to agriculture.

## Results

### A coarse-grained consumer-resource model under fluctuating environments

To determine whether environmental resource fluctuations alone could reproduce experimentally observed time series statistics, we considered a minimal consumer resource (CR) model in which *N*consumers compete for *M* resources via instantaneous growth dynamics described by

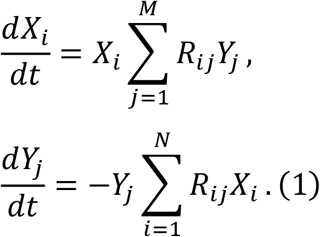

Here, *X*_*i*_ denotes the abundance of consumer *i, Y*_*j*_ the amount of resource *j*, and *R*_*ij*_ the consumption rate of resource *j* by consumer *i*. The resources in this model are defined at a coarse-grained level, such that individual resources could represent effective groups of metabolites or niches. We assumed that the resource consumption rates *R*_*ij*_ were independent of the external environment and constant over time, thereby specifying the intrinsic ecological properties of the community with a collection of *N* × *M* parameters.

To simplify this vast parameter space, we conjectured that the macroecological features of our experimental time series might be captured by a typical community consumption profile drawn from a larger statistical ensemble. Specifically, we considered an ensemble in which each *R*_*ij*_ was randomly selected from a uniform distribution between 0 and *R*_max_. To model the sparsity of resource competition within the community, each *R*_*ij*_ was set to zero with probability *S* (Fig. 1A). This ensemble approach allows us to represent arbitrarily large communities with a small number of global parameters, *S* and *R*_max_. Crucially, our ensemble assumes that there are no direct tradeoffs in resource use^4,5,19^, such that the maximum growth rate of each individual is strongly correlated to the total number of resources it can consume.

**Figure 1:**
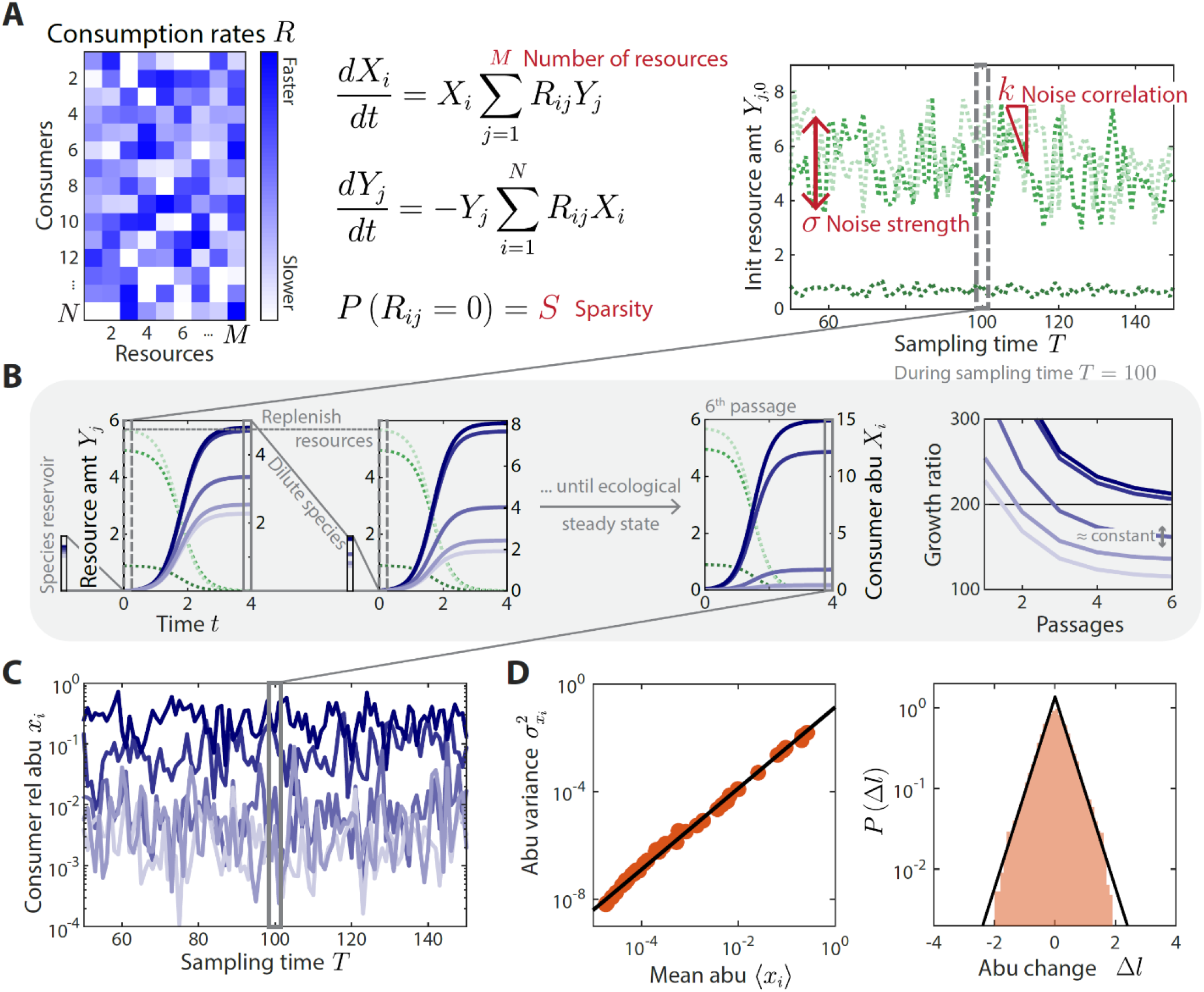
A coarse-grained consumer-resource model with fluctuating resource amounts. A) In the consumer-resource model, *X*_*i*_ denotes the abundance of consumer *i* and *Y*_*j*_ denotes the amount of coarse-grained resource *j*. The dynamics of the model are specified by consumption rates *R*_*ij*_ for *N* consumers and *M*resources. *R*_*ij*_ is drawn from a uniform distribution, and each *R*_*ij*_ is set to zero with probability *S*, the sparsity of resource competition. The initial resource amount *Y*_*j*,0_(*T*) at each sampling time *T* fluctuates with noise strength σ and restoring force *k*. *N* is estimated from each data set, and the 4 free ensemble-level parameters are highlighted in red. B) Shown are the dynamics of the model within one sampling time (*T* = 100, dashed gray box) for a subset of consumers and resources in a typical simulation. At each sampling time *T*, the model was simulated under a serial dilution scheme in which consumers (solid blue lines) grew until all resources (dotted green lines) were depleted, after which all consumer abundances were diluted by a fixed factor *D* = 200 and resource amounts were replenished to *Y*_0_(*T*). Each sampling time was initiated from an external reservoir of consumers, with all consumers present at equal abundance. Dilutions were repeated until an approximate ecological steady state was reached in which the ratios of final to initial abundances of all consumers changed by less than 5% of *D* between subsequent dilutions (Methods). The relative abundances at sampling time *T* were obtained from the final species abundances at steady state. C) The model maps a set of fluctuating resource amounts *Y*_*j*,0_(*T*) to a time series of species relative abundances *x*_*i*_(*T*) that can be compared to experimental measurements. D) The simulated time series in (C) exhibits statistical behaviors that reproduce those found in experiments, including a power-law scaling between the abundance variance and mean over time of each species (left) and an approximately exponential distribution of abundance changes (right). Black lines denote the best linear fit (left) and the best fit exponential distribution (right). The example simulation shown in (A-D) was generated with (*N, M, S*, σ, *k*) = (50, 30, 0.1, 0.2, 0.8).

We simulated the resource competition dynamics in Eq. 1 using a serial dilution scheme^20^ to mimic the punctuated turnover of gut microbiotas due to multiple feedings and defecations between sampling times. During a sampling interval *T*, each dilution cycle was seeded with an initial amount of each resource, *Y*_*j*,0_(*T*), and Eq. 1 was simulated until all resources were depleted (*dY*_*j*_/*dt* = 0 for all *j*). The entire community was then diluted by a factor *D* and resources were replenished to their initial amounts *Y*_*j*,0_(*T*) (Fig. 1B). To mimic the effects of internal or external reservoirs of species that could potentially compete for local resources^21^, we initialized the first dilution cycle of each sampling interval by assuming that *N* consumers were present at equal abundance. Additional dilution cycles were then performed until an approximate ecological steady state was reached (Fig. 1B, Methods). Consumer abundances at sampling time *T* were defined by this approximate ecological steady state. For most of the relevant parameter regimes we considered, this approximate steady state was reached within a reasonable number of generations (5-6 dilutions or ∼40 generations for *D* = 200). Our results did not depend on the precise composition of the reservoir (Fig. S1A), although they can strongly depend on the existence and the relative size of a reservoir (Fig. S1B). We discuss the implications of this finding in the Discussion.

Under the assumptions of this model, any temporal variation in consumer abundances must arise through external fluctuations in the initial resource levels *Y*_*j*,0_(*T*). To model these fluctuations, we assumed that the initial resource levels undergo a biased random walk around their average values 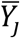:

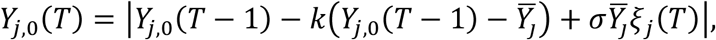

where *ξ*_*j*_(*T*) is a normally distributed random variable with zero mean and unit variance, σ determines the magnitude of resource fluctuations, and *k* is the strength of a restoring force that ensures the same resource environment on average over time (Fig. 1A). If *k* = 0, there is no restoring force and hence *Y*_*j*,0_(*T*) performs an unbiased random walk; if *k* = 1, *Y*_*j*,0_(*T*) fluctuates about its set point 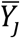 independent of its value at previous sampling time. As above, we used an ensemble approach to model the set points 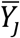, and assumed that each was independently drawn from a uniform distribution between 0 and *Y*_max_. This model yields a sequence of fluctuating resource amounts *Y*_*j*,0_(*T*), and a corresponding time series of consumer relative abundances *x*_*i*_(*T*) = *X*_*i*_(*T*)/ ∑_*k*_ *X*_*k*_ (*T*) (Fig. 1C).

The statistical properties of these time series are primarily determined by 5 global parameters: the total number of consumers in the reservoir *N*, the number of resources in the environment *M*, the sparsity of the resource consumption matrix *S*, and the resource fluctuation parameters σ and *k*. The absolute magnitudes of *R*_max_ and *Y*_max_ are not important for our purposes since they do not affect the predictions of consumer relative abundances. The precise value of the dilution factor and steady-state threshold did not substantially affect time series statistics, as long as a realistic number of generations elapsed between sampling times (Fig. S2). We extracted *N* from experimental data as the number of consumers that were present for at least one sampling time point, leaving only 4 free ensemble-level parameters. Previous studies have shown that the metabolic capabilities of bacterial species are more similar within families than between them^6,22,23^, thus we assumed that each consumer grouping *i* within our model represents a taxonomic family, and combined abundances of empirical operational taxonomic units (OTUs) or amplicon sequencing variants^24^ (ASVs) at the family level for analysis (Methods). Given the typical limits of detection of 16S amplicon sequencing data sets, consumers whose relative abundance was < 10^−4^ were set to zero abundance in both simulations and experimental data, and the abundances of the remaining consumers were normalized to a sum of one.

An example simulation using the parameters (*N, M, S*, σ, *k*) = (50, 30, 0.1, 0.2, 0.8) is shown in Fig. 1. This particular set of parameters produces relative abundance time series with highly similar statistical behaviors as in experiments involving daily sampling of human stool (Fig. 1D). Given these successes, we next systematically analyze the time series statistics generated by our model across parameter space, and compare against experimental behaviors to estimate model parameters for wide-ranging microbiotas.

### Model reproduces the statistics of human gut microbiota time series

To test whether our model can recapitulate all major features of experimental time series, we first focused on a data set of daily sampling of the gut microbiota from a human subject^9^ (Fig. 2). These data were previously shown^11^ to exhibit several distinctive statistical behaviors: 1) the variance 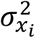 of family *i* over the sampling period scaled as a power law with its mean ⟨*x*_*i*_⟩ (Fig. 2B,F); 2) the log_10_(abundance change) Δ*l*_*i*_(*T*) = log_10_(*x*_*i*_(*T* + 1)/*x*_*i*_(*T*)), pooled over all families and across all sampling times, was well fit by an exponential distribution with standard deviation σ_Δ*l*_ (Fig. 2B,G); and 3) the distributions of residence times *t*_res_ and return times *t*_ret_ (the durations of sustained presence and absence, respectively) pooled over all families were well fit by power laws with an exponential cutoff (Fig. 2D,K). Through an exhaustive search of parameter space, we found that our model could reproduce all of these behaviors (Fig. 2F,G,K). In addition, several other important statistics were reproduced without any additional fitting: 1) the distribution of richness *α*(*T*), the number of consumers present at sampling time *T* (Fig. 2A,E), 2) the distribution of the restoring slopes *s*_*i*_ of the linear regression of Δ*l*_*i*_(*T*) against *l*_*i*_(*T*) ≡ log_10_(*x*_*i*_(*T*)) across all *T* (Fig. 2C,H), 3) the distribution of prevalences *p*_*i*_, the fraction of sampling times for which family *i* is present (Fig. 2A,I); 4) the relationship between *p*_*i*_ and ⟨*x*_*i*_⟩ (Fig. 2J); and 5) the rank distribution of mean abundance ⟨*x*_*i*_⟩ (Fig. 2L). Therefore, with only 4 parameters, our model was able to simultaneously capture at least 8 statistical behaviors in a microbiota time series, each of which may capture biologically relevant features of the community.

**Figure 2:**
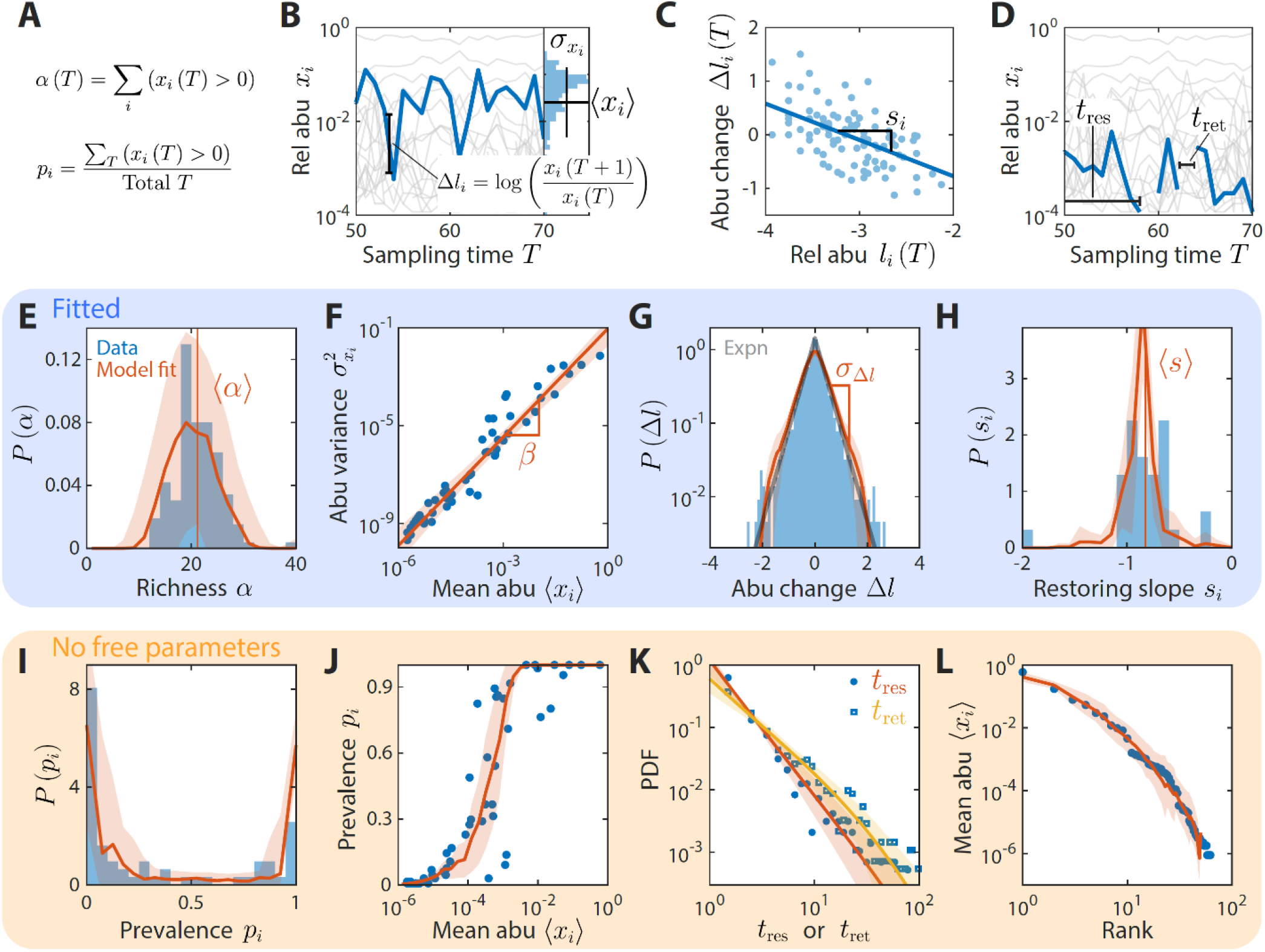
A coarse-grained consumer-resource model with fluctuating resource amounts reproduces experimentally observed statistics in the abundance time series from a daily time series of a human gut microbiota. In all panels, blue points and bars denote experimental data analyzed and aggregated at the family level^9^. Red lines and shading denote best fit model predictions as the mean and standard deviation, respectively, across 20 random instances of the best fit ensemble-level parameters, (*N, M, S, σ, k*) = (50, 30, 0.1, 0.2, 0.8). A-D) Illustrations of various time series statistics in (E-L). E) The distribution of richness *α*, the number of consumers present at a sampling time (A), and its mean ⟨*α*⟩ are well fit by the model. F) The variance 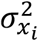 and mean ⟨*x*_*i*_⟩ over time (B) of each family’s abundance scale as a power law with exponent *β*. Here, *β* = 1.48 in experimental data and in simulations. G) The distribution of log_10_(abundance change) Δ*l* (B) across all families is well fit by an exponential with standard deviation *σ*_Δ*l*_. The gray line denotes the best fit exponential distribution, and is largely overlapping with the model prediction in red. H) The distribution of restoring slopes *s*_*i*_, defined based on the linear regression between the abundance change and the relative abundance for a species across time (C), is tightly distributed around a mean ⟨*s*⟩ that reflects the environmental restoring force. Best fit values of model parameters were determined by minimizing errors in ⟨*α*⟩, *β, σ*_Δ*l*_, and ⟨*s*⟩ (E-H, respectively). Using these values, our model also reproduced the distribution of prevalences (fraction of sampling times in which a consumer is present, D) (I), the relationship between prevalence and mean abundance (J), the distributions of residence and return times (durations of sustained presence or absence, respectively as illustrated in (D)) (K), and the rank distribution of abundances (L).

### Systematic characterization of the effects of consumer-resource dynamics on time series statistics

Since our model can reproduce the observed statistics in gut microbiota time series, we sought to determine how these statistics would respond to changes in model parameters, and thus how experimental measurements constrain the ensemble parameters across various data sets. To do so, we simulated our model across all relevant regions of parameter space. *S* and *k* were varied across their entire ranges, while *M* and *σ* were varied across relevant regions outside of which the model clearly disagreed with the observed data. For each set of parameters, each time series statistic was averaged across random instances of *R*_*ij*_ and *Y*_*j*,0_(*T*) drawn from the same statistical ensemble. For each statistic *z*, its global susceptibility *C*(*z, w*) to parameter *w* was calculated as the change in *z* when *w* is varied, averaged over all other parameters and normalized by the standard deviation of *z* across the entire parameter space. Due to the normalization, *C*(*z, w*) varies approximately between −3 and 3, where a magnitude close to 3 indicates that almost all the variance of *z* is due to changing *w*.

By clustering and ranking susceptibilities, we identified four statistics with |*C*(*z, w*)| > 2 that were largely determined by one of each of the four model parameters (Fig. 3, S3): mean richness ⟨*α*⟩, the power-law exponent *β* of 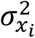 versus ⟨*x* ⟩, the standard deviation in log_10_(abundance change) *σ*_Δ*l*_, and the mean restoring slope ⟨*s*⟩ were almost exclusively susceptible to variations in *M, S, σ*, and *k*, respectively. Similar results were also obtained for local versions of the susceptibility, in which individual parameters were varied around the best fit values for the human gut microbiota in Fig. 2 (Fig. S4). Susceptibilities broadly illustrate how various time series statistics are affected by coarse-grained parameters of resource competition; we further investigate some specific examples in the next section.

**Figure 3:**
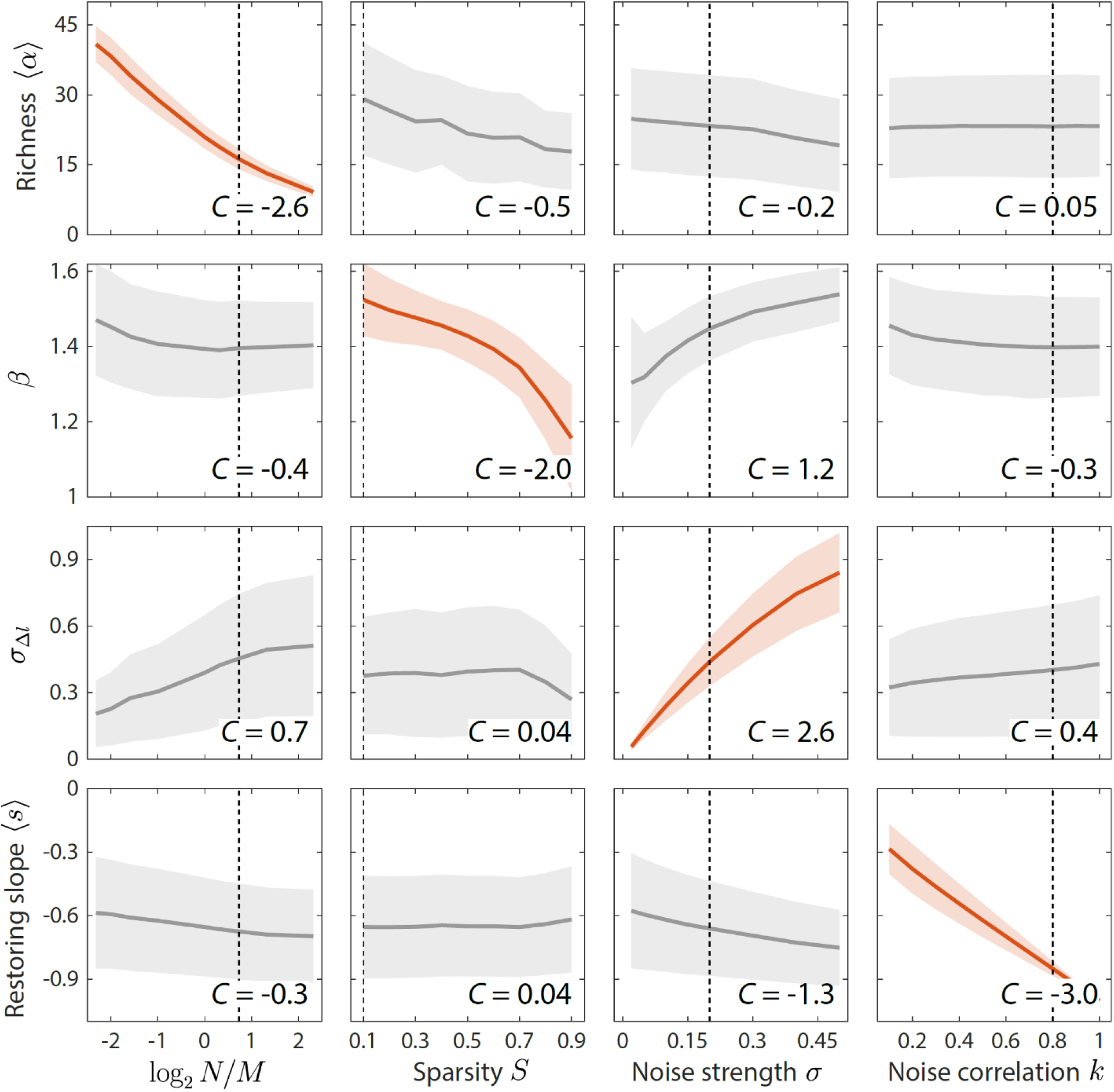
Macroscopic parameters of resource competition affect time series statistics in distinct manners. Shown are the changes in time series statistics (*y*-axis) in response to changes in model parameters (*x*-axis) for a comprehensive search across relevant regions of parameter space. Lines and shading show the mean and standard deviation of a statistic at the given parameter value across variations in all other parameters. Panels are highlighted in red when the corresponding susceptibility |*C*(*z, w*) > 2|, indicating that statistic *z* is strongly affected by parameter *w* regardless of the values of other parameters. Dashed lines highlight best fit parameter values to the experimental data in Fig. 2. Simulations were carried out for *N* = 50 across *M*∈ [10, 20, 30, 40, 50, 100, 150, 200, 250], *S* ∈ [0.1, 0.9] in increments, σ ∈ [0.05, 0.5] in 0.05 increments, and *k* ∈ [0.1, 1] in 0.1 increments.

The exclusive susceptibilities of these four statistics suggest that they can serve as informative metrics for estimating model parameters. Therefore, we estimated model parameters by minimizing the sum of errors between model predictions and experimental measurements of these four statistics, and obtained estimation bounds by determining parameter variations that would increase model error by 5% of the mean error across all parameter space. As we will show, the resulting bounds are small relative to the differences among distinct microbiotas, indicating that meaningful conclusions can be drawn from the best-fit values of the ensemble-level parameters of resource competition. In summary, the 4 model parameters were fit to 4 summary statistics, mean richness ⟨*α*⟩, variance-mean scaling exponent *β*, standard deviation of abundance change *σ*_Δ*l*_, and mean restoring slope ⟨*s*⟩ (Fig. 2E-H, respectively). The shape of their corresponding distributions and scalings, as well as at least 4 other statistics, are all parameter-free predictions (Fig. 2I-L), providing support for our model.

### Origins of distinctive statistical behaviors in species abundance time series

To understand the mechanisms that underlie the susceptibilities of various time series statistics to model parameters, we investigated their origins within our model, focusing on how they constrain the parameters.

The average richness ⟨*α*⟩ is a fundamental descriptor of community diversity. Within our model, ⟨*α*⟩ is largely determined by and increases with increasing resource number *M*(*C*(*α, N*/*M*) = −2.6), as expected for CR dynamics. The sparsity of resource use *S* impacts the power-law exponent *β* between 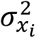 and ⟨*x* ⟩ (*C*(*β, S*) = −2.0). The effect of *S* on *β* can be partially understood as follows. When sparsity is high (*S* ≈ 1), consumers consume distinct sets of resources with little competition from other consumers, and the variation in the number of utilized resources can be large relative to the average value *M*(1 − *S*). In this case, both 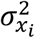 and ⟨*x* ⟩ scale with the number of resources consumed, and hence *β* ≈ 1. By contrast, when sparsity is low (*S* ≈ 0), all consumers consume all resources and hence the numbers of resources consumed are the same among consumers. To better understand this limit, we simulated a no-competition model in which all consumers consume distinct sets of the same number of resources. For large number of resources, these simulations predicted that *β* ≈ 1.5, in agreement with our model for *S* ≈ 0 (Fig. S5). Simulations of the no-competition model with varying numbers of resources consumed predicted values of *β* between 1 and 1.5 (Fig. S5), suggesting that the effect of *S* on *β* can be partially attributed to differences in the number of resources consumed across consumers. Together, *α* and *β* constrain the parameters of resource competition *M*and *S*.

The distribution of Δ*l* describes the nature of abundance changes. As expected, the width of the distribution is largely determined by and increases with increasing *σ* (*C*(*σ*_Δ*l*_, *σ*) = 2.6). For the gut microbiota data set in Fig. 2, the shape of the distribution was well fit by an exponential. Within our model, the shape of the distribution aggregated across all consumers is determined by *N*/*M*and the sparsity *S*, emerging from the mixture of each consumer’s individual distribution (Fig. S6). When *N*/*M*< 1 and the sparsity *S* is low, individual distributions of Δ*l* are well fit by normal distributions, and pool together to generate another normal distribution. When *N*/*M*< 1 and sparsity *S* is high, individual distributions remain normal, but can pool together to generate a non-normal distribution that is well fit by an exponential (see also Ref. ^25^). By contrast, when *N*/*M* > 1, individual distributions can be well fit by an exponential, and can pool together to approximate another exponential. Simulations of the no-competition model considered above led to individual and aggregate distributions that were normal in all cases, indicating that resource competition is responsible for generating the non-normal distributions of Δ*l* in our model (Fig. S5). Although it is challenging to discern the shape of individual distributions in most experimental data sets given the limited numbers of samples, the shape of the aggregate distribution of Δ*l* informs the parameters of resource competition *M* and *S*. In particular, an exponential distribution of Δ*l* suggests either significant resource competition in the form of *N* > *M*, or substantial niche differentiation in the form of high *S*. Other statistics such as *β* can help to distinguish between these two regimes.

The distribution of restoring slopes *s*_*i*_ describes the tendency with which consumers revert to their mean abundances following fluctuations. As expected, the mean ⟨*s*⟩ is almost completely determined by *k*, which describes the autocorrelation in resource levels (−⟨*s*⟩ ≈ *k* and *C*(⟨*s*⟩, *k*) = −3.0). Together, the distributions of Δ*l* and *s*_*i*_ constrain the parameters of external fluctuations *σ* and *k*.

Within our model, resource fluctuations can lead to the temporary “extinction” of certain species when they drop below our detectability threshold *x*_*i*_(*T*) < 10^−4^. The distributions of residence and return times, *t*_res_ and *t*_ret_, reflect the probabilities of extinction as well as correlations between sampling times. For all parameter sets explored, these distributions can be well fit by power laws, with an exponential cutoff to account for finite sampling in time^11^. As expected, the power-law slopes *v*_res_ and *v*_ret_ decrease (become more negative) with increasing *σ* or *k*, since increasing external noise or decreasing correlations in time increases the probability of fluctuating between existence and extinction for each consumer, thereby decreasing *v*_res_ and *v*_ret_ for all consumers. By contrast, *v*_res_ and *v*_ret_ change in opposite directions in response to variation in *M*. Increasing *M*leads to a larger number of highly prevalent consumers, thereby increasing the mean and broadening the distribution of *t*_res_ and decreasing the mean and narrowing the distribution of *t*_ret_. Since the 4 ensemble-level parameters are already fixed by other statistics, the distributions of *t*_res_, *t*_ret_, and *p*_*i*_ are parameter-free predictions of our model. In other words, a macroscopic characterization of the effective resource competition and resource fluctuations is sufficient to predict the statistics of “extinction” dynamics, as well as the abundance rank distribution and the relationship between consumer abundance and prevalence.

Having clarified the origins of several important time series statistics, we next investigated how they depend on model assumptions. First, we considered a non-interacting null model in which consumer abundances were drawn from independent normal distributions whose means and variances were fitted directly from the data. Even with a large number of free parameters, this null model was unable to capture some of the time series statistics reproduced by our resource competition model above, such as the distributions of residence and return times (Fig. S7). We reasoned that the discrepancies between experimental data and the null model could be due to the lack of interspecies interactions. To test this hypothesis, we examined the pairwise correlations between the abundances of pairs of consumers across sampling times. The measured distribution of pairwise correlations is much broader than the prediction of the non-interacting model, which is sharply peaked about zero as expected (Fig. 4). By contrast, the distribution of correlations predicted by our resource competition model was in much closer agreement with the experimental data without any additional fitting parameters (Fig. 4). These findings suggest that competitive interactions among consumers are required to capture important details of community dynamics.

**Figure 4:**
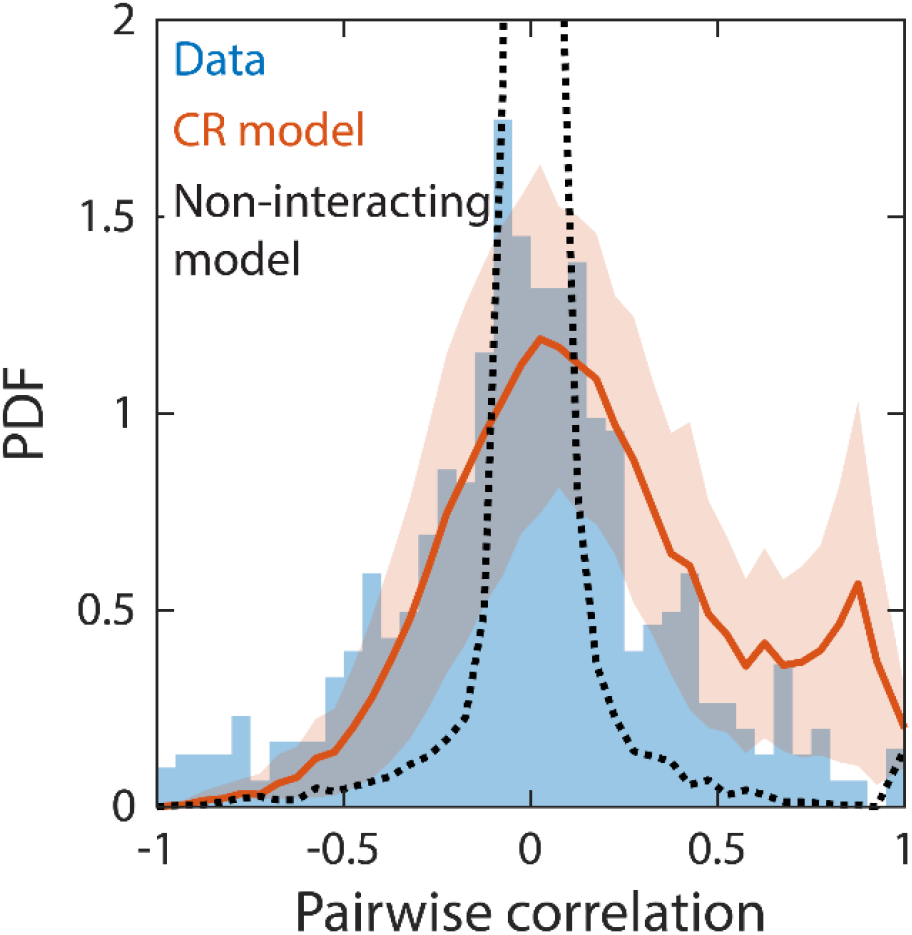
Correlations between abundances of consumer pairs were captured by the consumer-resource model, but not by a non-interacting null model. Shown in blue is the distribution of correlations between the abundances across sampling times of all consumer pairs for the experimental data in Fig. 2. Red line represents parameter-free model predictions as in Fig. 2, using the same best fit parameters; shading represents 1 standard deviation. Black dashed line shows predictions of a non-interacting null model in which consumer abundances were drawn from independent normal distributions whose mean and variance were extracted from data.

Second, since the distributions of Δ*l, t*_res_, and *t*_ret_ are dependent on correlations between sampling times, it was initially puzzling that their distributions in some data sets remained similar after shuffling sampling times, raising doubt as to what extent these statistics hold information about the underlying intrinsic dynamics^26^. Our results reconcile the apparent conundrum, since within our model richness ⟨*α*⟩ and Taylor’s law exponent *β* do not depend on correlations between sampling times and are also the statistics that are most informative about the intrinsic parameters *M*and *S* (Fig. 3). As a result, the shuffled time series were well fit by our model, and yielded best fit values that were identical to those produced by the actual time series but with *k* = 1, as expected due to the absence of correlation across sampling times (Fig. S8). Moreover, we found that many microbiotas yielded best fit values of *k* close to 1, indicating low correlation across sampling times (Fig. 5). Taken together, our results demonstrate that although external fluctuations in resource levels are responsible for generating species abundance variations, the intrinsic properties of resource competition determine the resulting scaling exponents of the statistical behaviors.

**Figure 5:**
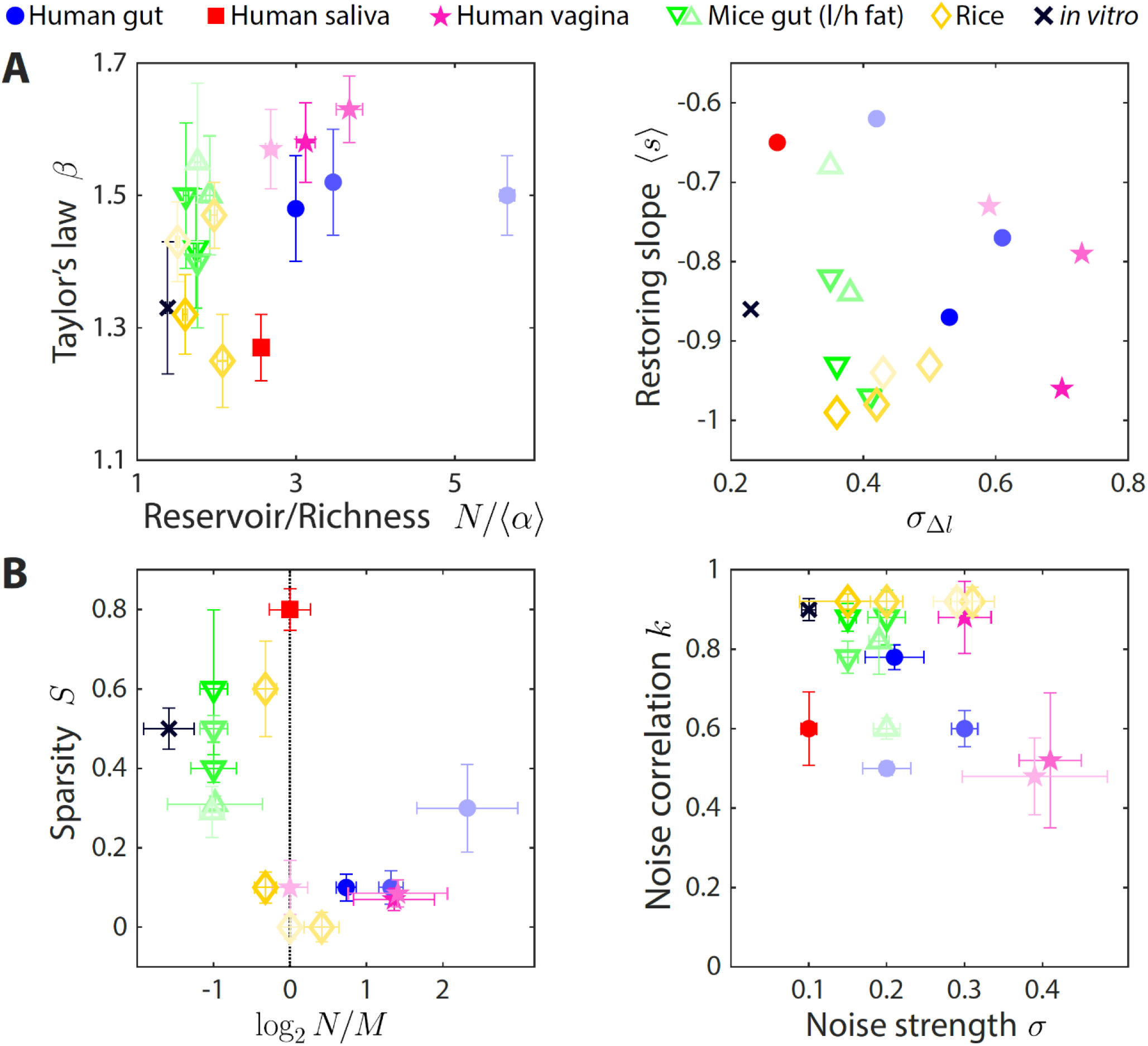
The statistics of wide-ranging microbiotas were captured by the coarse-grained consumer-resource model in different regimes of resource competition and environmental fluctuations. Shown are time series statistics (A) and corresponding best fit model parameters (B) for human microbiotas from stool^9,10^ (blue circles), saliva^10^ (red square), and the vagina^28^ (pink stars), gut microbiotas of mice under low-fat (green downward triangles) and high-fat (green upward triangles) diets^27^, and plant microbiotas from the rice endosphere, rhizosphere, rhizoplane, and bulk soil^29^ (diamonds). A) Microbiota origin generally dictates the scaling exponent *β* and the ratio between the reservoir size *N* (number of observed families throughout the time series) and the richness ⟨*α*⟩ (left), as well as the mean restoring slope ⟨*s*⟩ and standard deviation of log_10_(abundance change) (right). Error bars denote 95% confidence intervals. B) Microbiota origin generally dictates the best fit parameters of resource competition, *N*/*M*and *S* (left), and of environmental fluctuations, *σ* and *k* (right). Error bars denote variation in the parameter that would increase model error (as interpolated between parameter values scanned) by 5% of the mean error across all parameter values scanned.

Lastly, grouping time series at coarser taxonomic levels yielded similar qualitative statistical behaviors, which were also reproduced by our model (Fig. S9). Interestingly, the best fit value of *S* was larger for coarser groupings, supporting the hypothesis that higher taxonomic groups have substantially differentiated metabolic capabilities.

Taken together, our analyses demonstrate the complex relationships among time series statistics and highlight the unification of these statistics within our model using only a small number of biologically meaningful parameters.

### Time series statistics distinguish wide-ranging microbiotas

Having developed a simple method to estimate model parameters, we applied this method to time series of wide-ranging microbial communities. In addition to microbiotas from the human and mouse gut^9,10,27^, we examined communities from the human vagina^28^, human saliva^10^, and in and around rice roots^29^. Their time series statistics varied broadly in value (Fig. 5A). Nevertheless, our model successfully reproduced the experimental statistics across all communities (Fig. S10-S15), suggesting that simple resource competition models can capture many of the macroscopic features of these microbiotas.

The best fit parameters suggest that the effective resource competition dynamics occur in distinct regimes (Fig. 5B). Human gut microbiotas were best described by *N* > *M*, suggesting that there are more species in the reservoir than resources in the environment, by contrast to mouse gut microbiotas that were best described by *N* < *M*. In terms of resource niche overlaps, human gut microbiotas were best fit with sparsity *S* < 0.3, while mouse gut microbiotas were best fit with *S* > 0.3, indicating that on average, pairs of bacterial families are more metabolically distinct in the mouse versus the human gut.

Unlike gut microbiotas, a human saliva microbiota yielded best fit parameters *N* ≈ *M* and *S* ≈ 0.8, suggesting that this community has access to abundant resources and that each effective resource is competed for by a small fraction of the extant bacterial families. All vaginal microbiotas were best fit with *S* < 0.1, suggesting intense resource competition. Interestingly, although vaginal microbiotas can be classified into several types based on the dominating species^28^, time series statistics among the different types (and hence their best fit parameters) were virtually indistinguishable (Fig. 5B, S12).

Like vaginal microbiotas, microbial communities residing in the bulk soil around rice roots and in the associated rhizoplane and rhizosphere were well described by *S* < 0.1. By contrast, the community in the associated endosphere was best described by *S* ≈ 0.6, suggesting that resource competition is less fierce within plant roots than around them. However, we caution that the stable point of these plant-associated communities shifts over time^29^, which may affect time series statistics outside the scope of our model.

In addition, inferences about the nature of environmental fluctuations can be made from the best fit values of *σ* and *k* (Fig. 5B). Across all but two vaginal microbiota data sets, *σ* ≈ 0.2 ± 0.1, indicating that changes in resource levels smaller than this magnitude will generate abundance changes that look like typical fluctuations. The best fit values of *k* varied between 0.5 and 1 across data sets, suggesting that the dynamics of microbial communities occur faster than or comparable to the typical sampling frequency of longitudinal studies.

Inferences about intrinsic parameters of resource competition and external parameters of environmental fluctuations were also consistent with expectations for *in vitro* passaging of a complex community derived from humanized mice^30^. The resulting time series statistics were best fit by the smallest value of *σ* among the data sets studied, indicating that the *in vitro* environment has low noise across sampling times; the nonzero *σ* presumably arises from technical variations that result in effective noise in resource levels. The best fit value of *M* was larger than the reservoir size *N*, suggesting that there are many distinct resources in the complex medium used for passaging and consistent with the ability of more diverse inocula to support more diverse *in vitro* communities^30^. The consistency of these results further supports the mechanistic picture provided by our modeling framework. Taken together, our model predicts ensemble-level parameters of resource competition and external parameters of environmental fluctuations for several widely studied microbial communities that can inform future mechanistic studies.

## Discussion

Here, we presented a coarse-grained CR model that generates species abundance time series from fluctuating environmental resources. We demonstrated that this model reproduces several important statistical behaviors, and how these observations constrain the parameters of resource competition within the model. Moreover, we successfully fitted the model to wide-ranging microbiotas, which allowed us to draw conclusions about their parameters of effective resource competition. In sum, our work provides a framework that unifies a plethora of time series statistics and exploits them to learn about the underlying community dynamics.

An important and novel property of our model is its ability to reproduce many statistical behaviors across wide-ranging data sets with a minimal number of effective parameters. To what extent these effective parameters can be interpreted mechanistically should be an exciting avenue of future investigation. Although by no means exhaustive, our framework nevertheless addresses several pertinent questions regarding construction of useful models of microbiota dynamics. The success of our model in reproducing experimental time series statistics is consistent with bioinformatics-guided analyses of complex communities demonstrating that metabolic capability is a major determinant of community composition^22,23^. Our results also suggest that the contributions of a persistent reservoir of species are important for the dynamics of wide-ranging microbiotas^21^. Within our model, the lack of a reservoir renders poor consumers unable to recover to meaningful abundance within a sampling time even when resource fluctuations are in their favor, thereby distorting time series statistics. Further experimental work is required to ascertain the amount of growth and change that occurs during sampling time scales.

In terms of intrinsic metabolic properties, our results provide a baseline expectation for the effective number of resources or available niches in the wide-ranging systems examined here, and to what extent they are competed for by extant consumers. In terms of environmental properties, our results provide a baseline expectation to help distinguish between typical fluctuations and large perturbations in resources. These expectations can aid in the engineering of complex microbiotas.

In general, our work demonstrates that it is feasible to reproduce time series statistics using consumer-resource models of microbiota dynamics, thereby generating mechanistic hypotheses for further investigation. In future work, more detailed hypotheses can be generated by investigating how time series statistics are affected by modifications to baseline CR dynamics, such as the incorporation of metabolic cross-feeding^6,31^, functional differentiation from genomic analysis^32-34^, and physical variables such as pH^35,36^, temperature^37^, and osmolality^38^. In addition, recent studies have shown that evolution can substantially affect the dynamics of human gut microbiotas^39-41^. It will therefore be illuminating to incorporate evolutionary dynamics into CR models under fluctuating environments^19^. Such extended models can then be applied to probe the underlying mechanisms in microbiotas with urgent applications, including those in marine environments, wastewater treatment plants, and the guts of insect pests and livestock.

## Methods

### Simulations of a CR model with fluctuating resource amounts

Under a serial dilution scheme, an ecological steady state is reached when the dynamics in subsequent passages are identical, which is the case when all consumers are either extinct or have a growth ratio (the ratio of a consumer’s final and initial abundances within one passage) equal to the dilution factor *D*. Due to the slow path to extinction of some consumers, reaching an exact ecological steady state can require hundreds of passages, presumably more than realistically occurs between sampling times in the data sets examined here. We assumed instead that between sampling times an ecological steady state is only approximately reached, defined as the growth ratios of all species changing by less than a threshold fraction of *D* between subsequent passages. Throughout this study, *D* was set to 200 and the steady state threshold was 5%, under which a steady state was approximately reached in about 5 dilutions (Fig. 1B). In this manner, our model assigns a well-defined state of consumer abundances to each resource environment while ensuring that only a reasonable amount of change occurs between sampling times. Note that in human gut microbiotas, abundances can change by more than 1,000 fold between daily samplings, indicating that at least 10 generations elapsed between sampling times (Fig. 2B). The precise value of *D* did not affect time series statistics, and steady state thresholds between 1% and 10% generated similar time series statistics (Fig. S2). We therefore expect our results to hold regardless of the precise values of these two parameters. Simulations were carried out in MATLAB, and all codes are freely available online at https://bitbucket.org/kchuanglab/consumer-resource-model-for-microbiota-fluctuations/.

### Analysis of 16S amplicon sequencing data

Raw 16S sequencing data from Refs. ^10,28^ were downloaded from SRA, and ASVs were extracted using DADA2^24^ with default parameters. OTUs or ASVs from other studies were downloaded and analyzed in their available form.

## Acknowledgements

We thank members of the Huang lab and Lisa Maier, Rui Fang, Jie Lin, and Felix Wong for helpful discussions. We thank Stephanie Song and Nicholas Chia for sharing metadata. This work was funded by a Stanford School of Medicine Dean’s Postdoctoral Fellowship (to P.H.), a Stanford Terman Fellowship (to B.H.G.), and NIH Awards R01 AI147023 and RM1 GM135102 (to K.C.H.). K.C.H. is a Chan Zuckerberg Biohub Investigator.

## Supplemental figures

**Figure S1:**
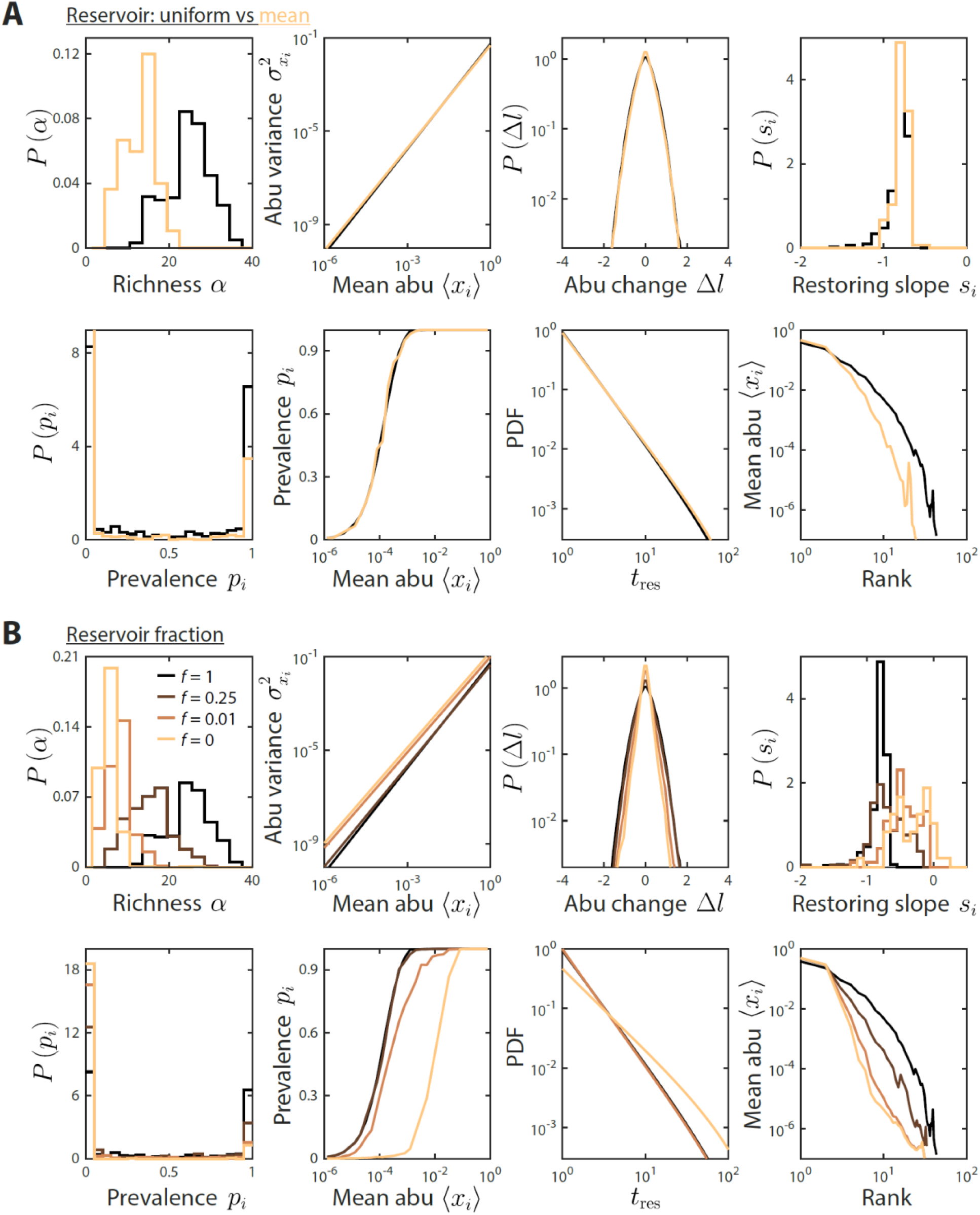
Reservoir composition does not substantially affect time series statistics. A) Depicted are mean model predictions using the parameter set in Fig. 1 and 2, with two definitions of the reservoir of consumers used to initialize the dynamics at each sampling time: a uniform reservoir in which all consumers are present at equal abundance (black), and a mean reservoir equal to the steady state composition given by the set point environment 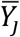 initialized by a uniform reservoir (light brown). Most statistics were not substantially affected. The richness was lower when initializing with the mean reservoir, but the susceptibility of richness to model parameters is expected to remain qualitatively the same. B) Depicted are mean model predictions with varying reservoir fraction *f*, in which initial consumer abundances during sampling time *T* were determined by a combination of the reservoir and the steady state at the previous sampling time, 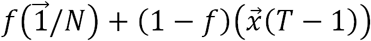. Many statistics were substantially affected by the value of *f*. Notably, the contribution from the previous steady state introduces autocorrelations, thereby increasing the mean restoring slope. Moreover, the absence of a reservoir substantially decreased richness since low-abundance consumers cannot grow enough even when resource levels fluctuate in their favor.

**Figure S2:**
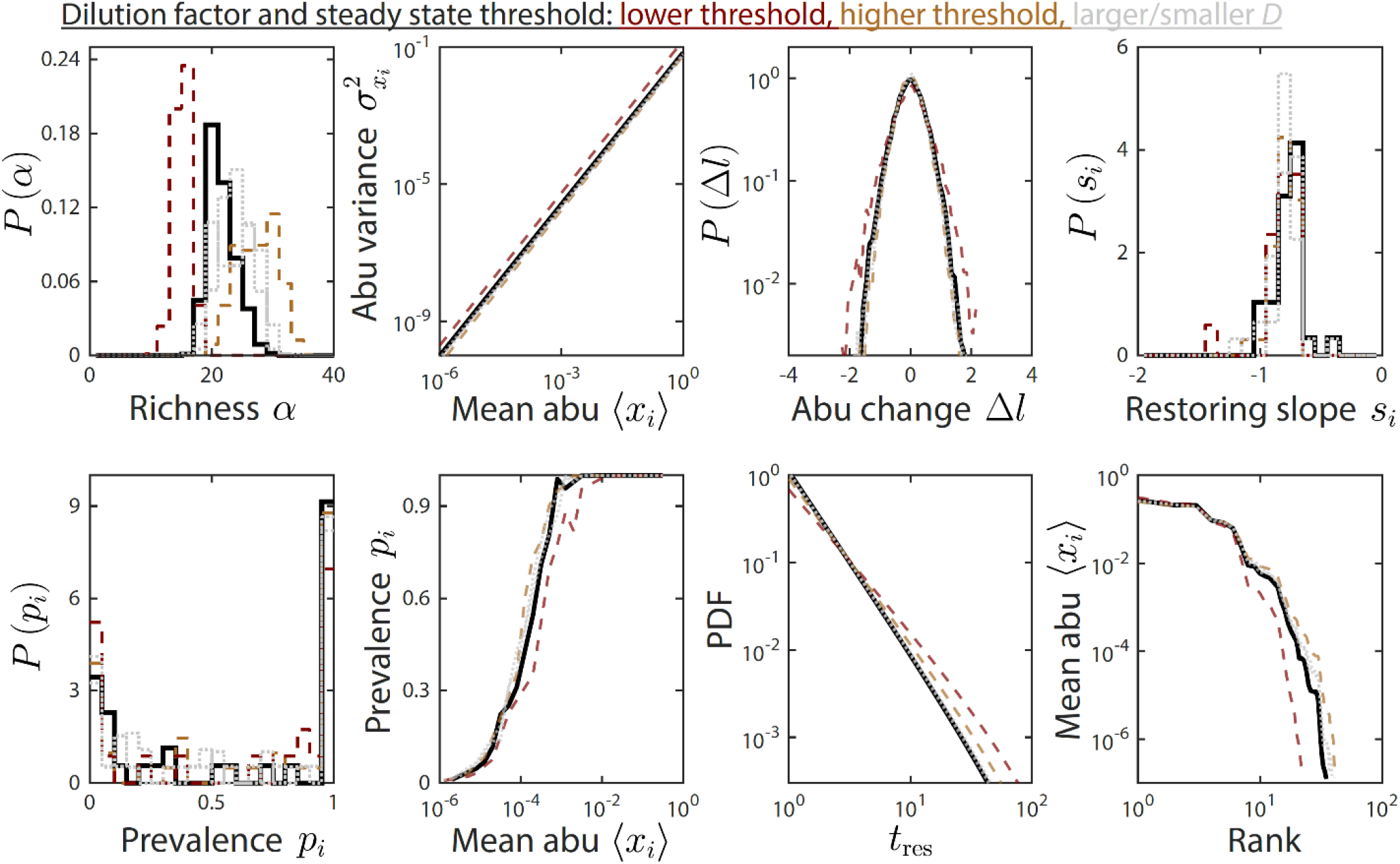
The dilution factor and steady state threshold do not substantially affect time series statistics. Depicted are time series statistics from one instance of the parameter set used in Figs. 1 and 2, simulated using a dilution factor *D* = 200 and a steady state threshold of 5% (solid black line), *D* = 40 or *D* = 1,000 (dotted gray lines), and a threshold of 1% and 10% (dashed brown and orange lines, respectively).

**Figure S3:**
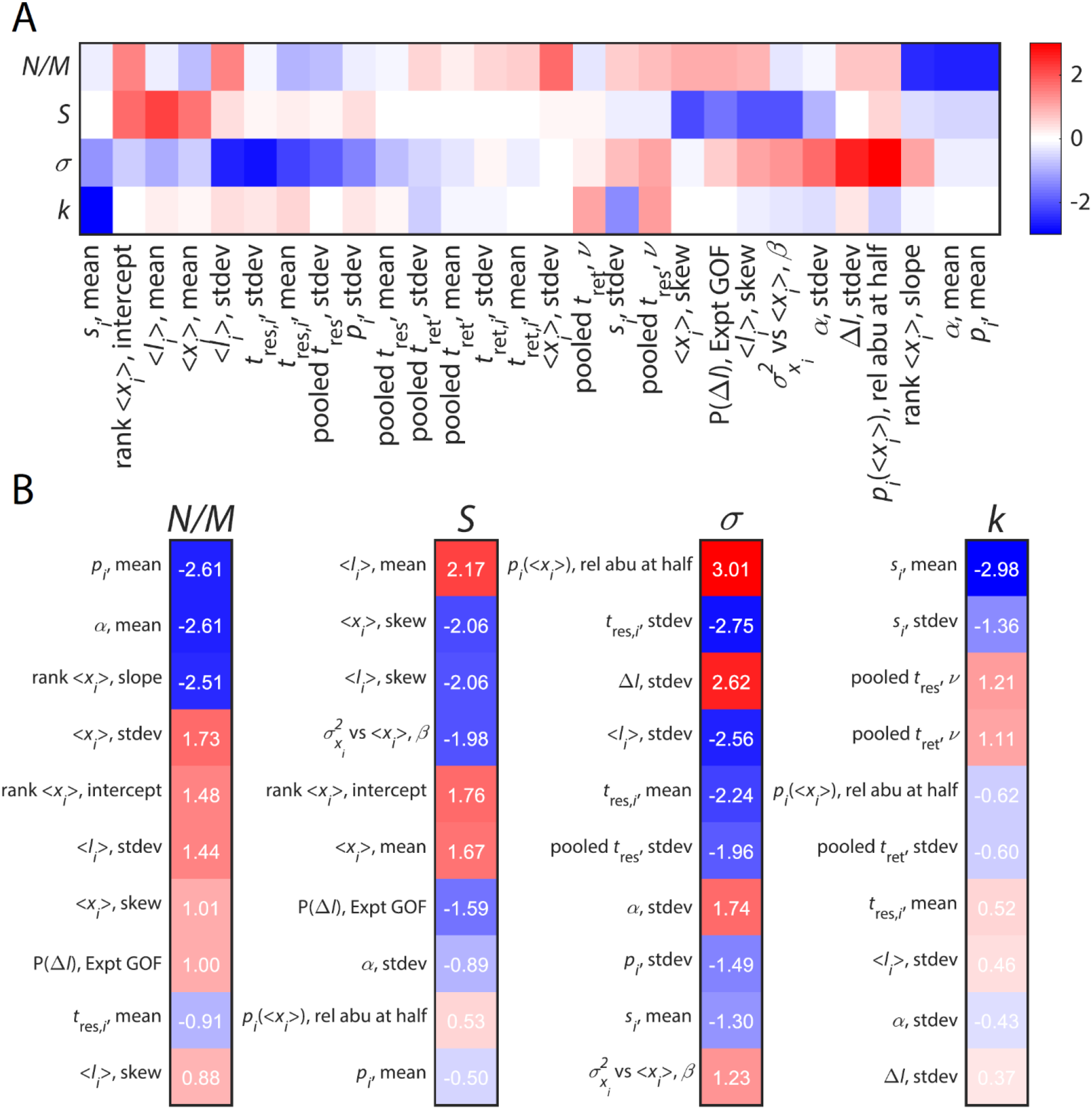
Time series statistics are differentially susceptible to model parameters. 28 time series statistics were clustered (A) and ranked (B) according to their susceptibilities to identify statistics that strongly constrain the values of key parameters.

**Figure S4:**
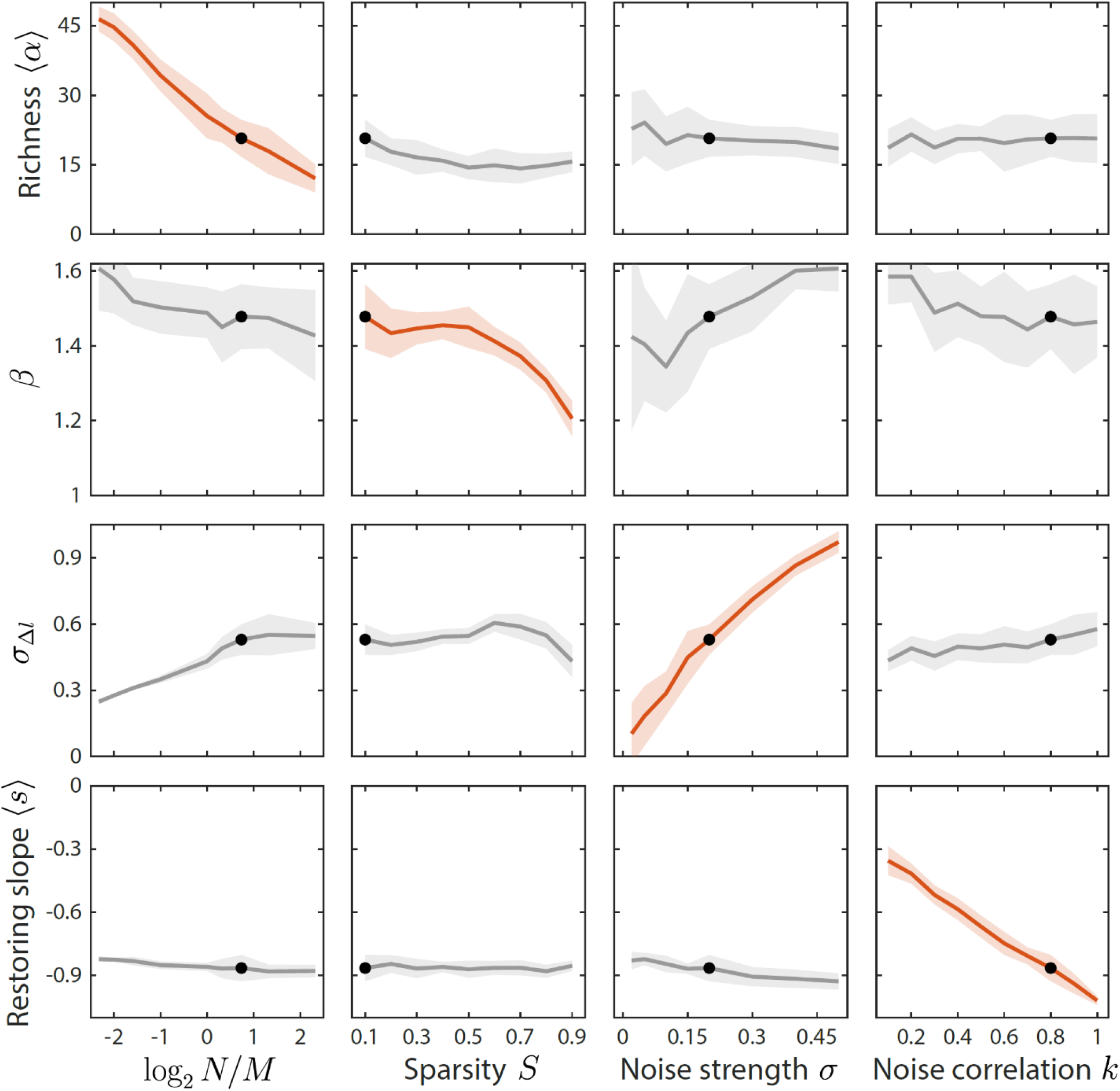
Local susceptibilities behave similarly to their global counterparts. Shown are changes in time series statistics (*y*-axis) in response to changes in model parameters (*x*-axis) as in Fig. 3. Here, lines and shading show the mean and standard deviation of a statistic at the given parameter value with other parameters fixed to their best-fit values. Black dots denote the best-fit values.

**Figure S5:**
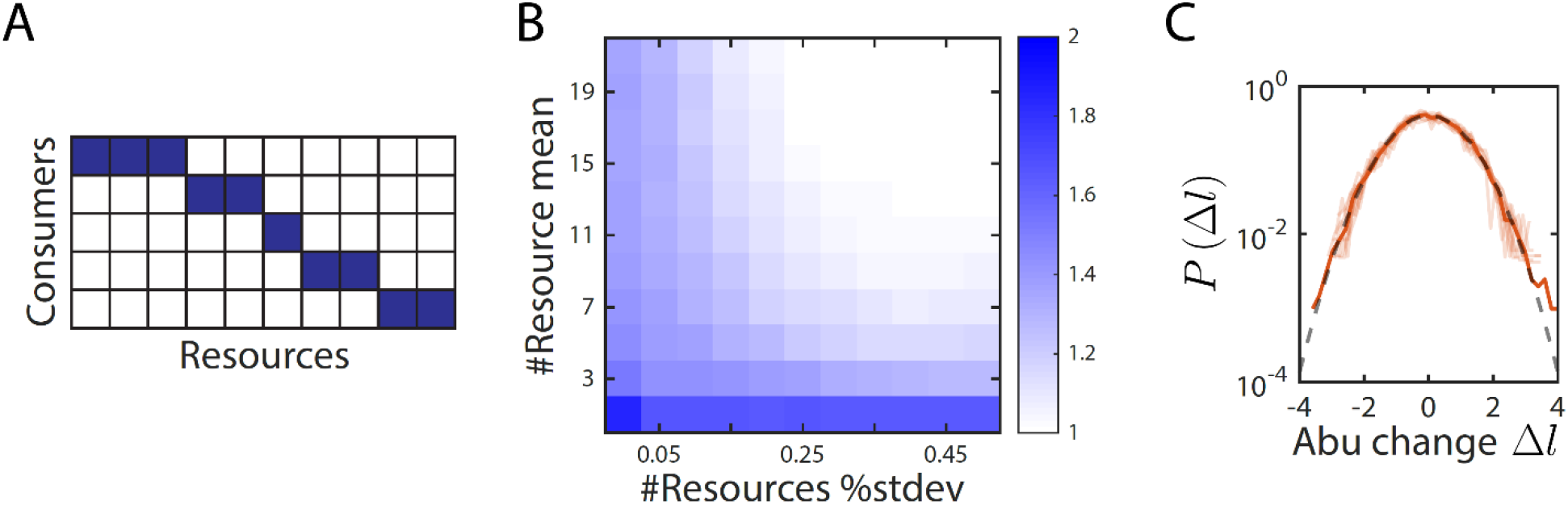
The no-competition model provides an intuitive explanation of variations in the scaling exponent *β* between 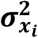 and ⟨*x*_*i*_⟩. A) In the no-competition model, each consumer consumes a disjoint set of resources. B) The mean and standard deviation of the number of resources consumed per consumer determine the scaling exponent *β*. Shown is the average value of *β* over 1,000 random instances of the no-competition model, across values of the mean number of resources consumed per consumer (*y*-axis) and the standard deviation in the number of resources consumed divided by the mean (*x*-axis). In this example, the model involved 10 consumers. At high variance in the number of resources consumed, *β* → 1, whereas at low variance, *β* varies between 1 and ≈ 1.5 depending on the mean number of resources consumed. C) Aggregate (solid orange line) and individual (light orange lines) distributions of abundance changes are normal (grey dashed line) in the no-competition model. The distributions of abundance changes, normalized by their sample standard deviations, are shown for an example simulation with 7±1 resources per consumer.

**Figure S6:**
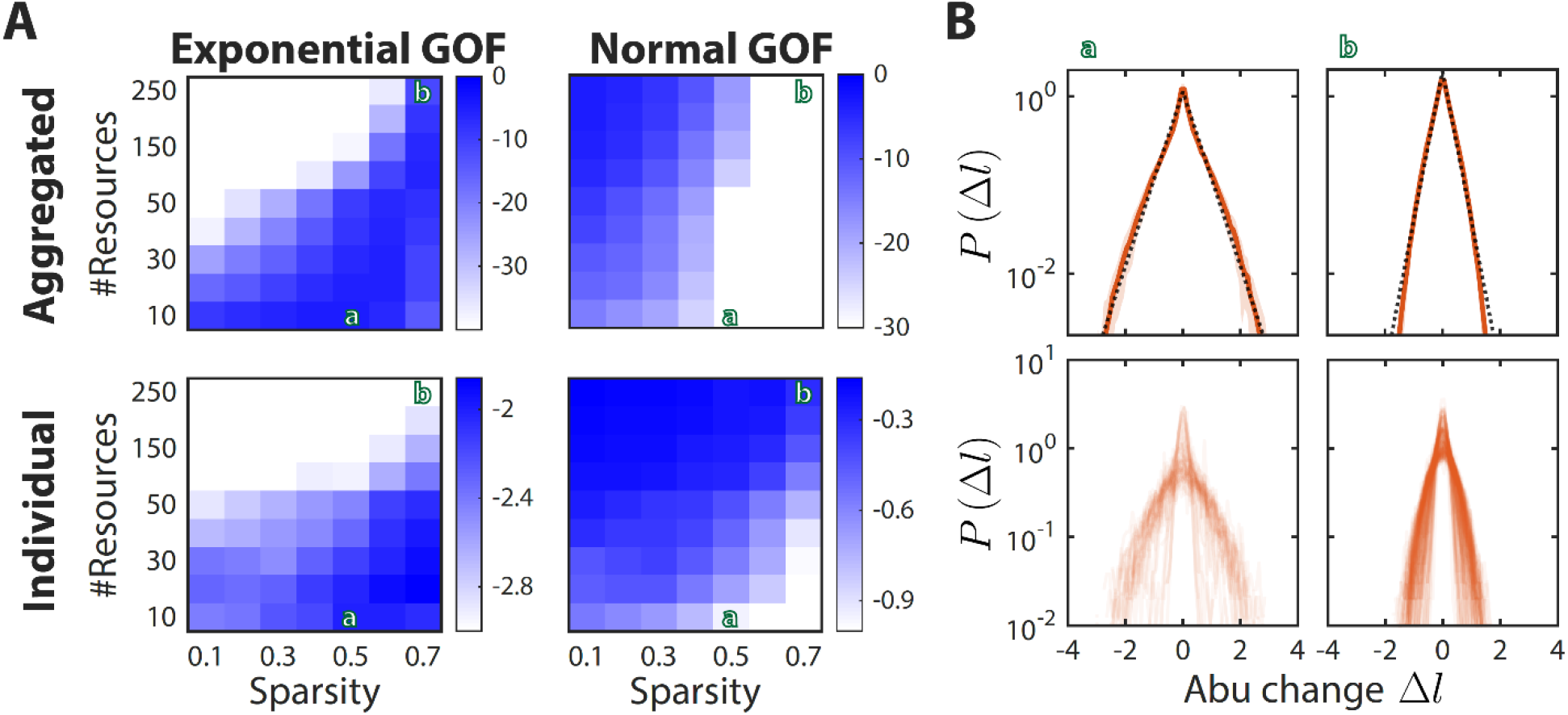
Sparsity and the number of metabolites determine the shape of the distribution of abundance changes *P*(*Δl*). A) Shown is the average goodness of fit (as determined by the *p*-value of the Kolmogorov-Smirnov test) of *P*(*Δl*) to an exponential (left) or a normal (right) distribution, for the distribution aggregated over all consumers (top) or the median value across the distributions of individual consumers (bottom). Larger values (blue) denote better fits. Green letters denote examples shown in (B). The other model parameters were fixed at (*N, σ, k*) = (50, 0.2, 0.8) as in Fig. 2. B) Shown are two different regimes that result in an exponential distribution of abundance changes. Example (a) demonstrates that when *N*/*M*> 1, the aggregated distribution is better fit by an exponential (black dotted line) than by a normal distribution (top), and that the median of the individual distributions is also better fit by an exponential (bottom). Example (b) demonstrates that when *N*/*M*< 1 and *S* is high, the aggregated distribution is still better fit by an exponential, but the median of the individual distributions is better fit by a normal distribution.

**Figure S7:**
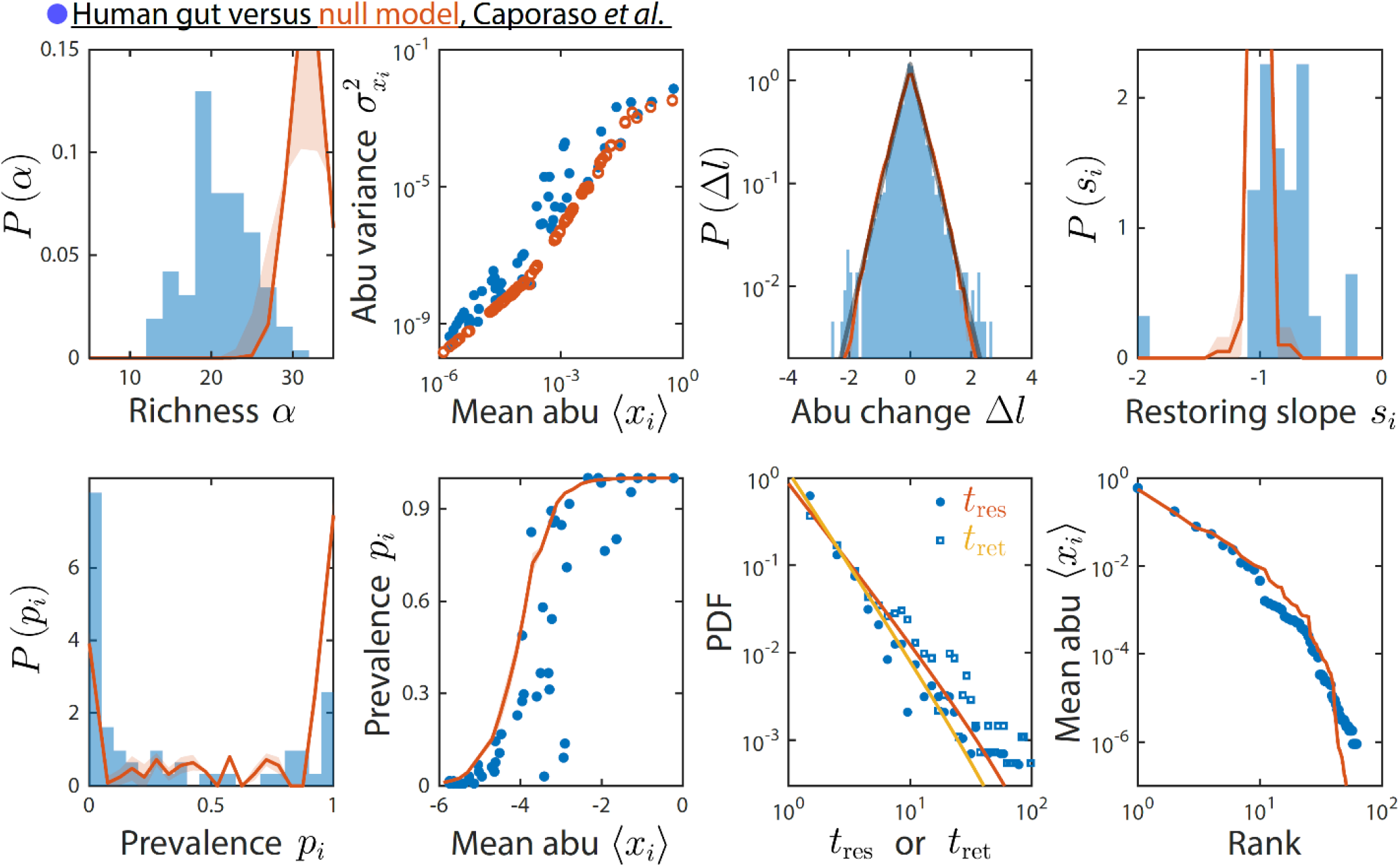
A non-interacting null model reproduced some, but not all, time series statistics. Shown are data from Ref. ^9^ as in Fig. 2 and comparisons to a non-interacting null model in which consumer abundances were drawn from independent normal distributions whose mean and variance were extracted from data. As a result of its many inputs, the null model was able to reproduce statistics such as the rank distribution of abundances, but was unable to reproduce the distributions of residence and return time, nor the distribution of pairwise correlations (Fig. 4).

**Figure S8:**
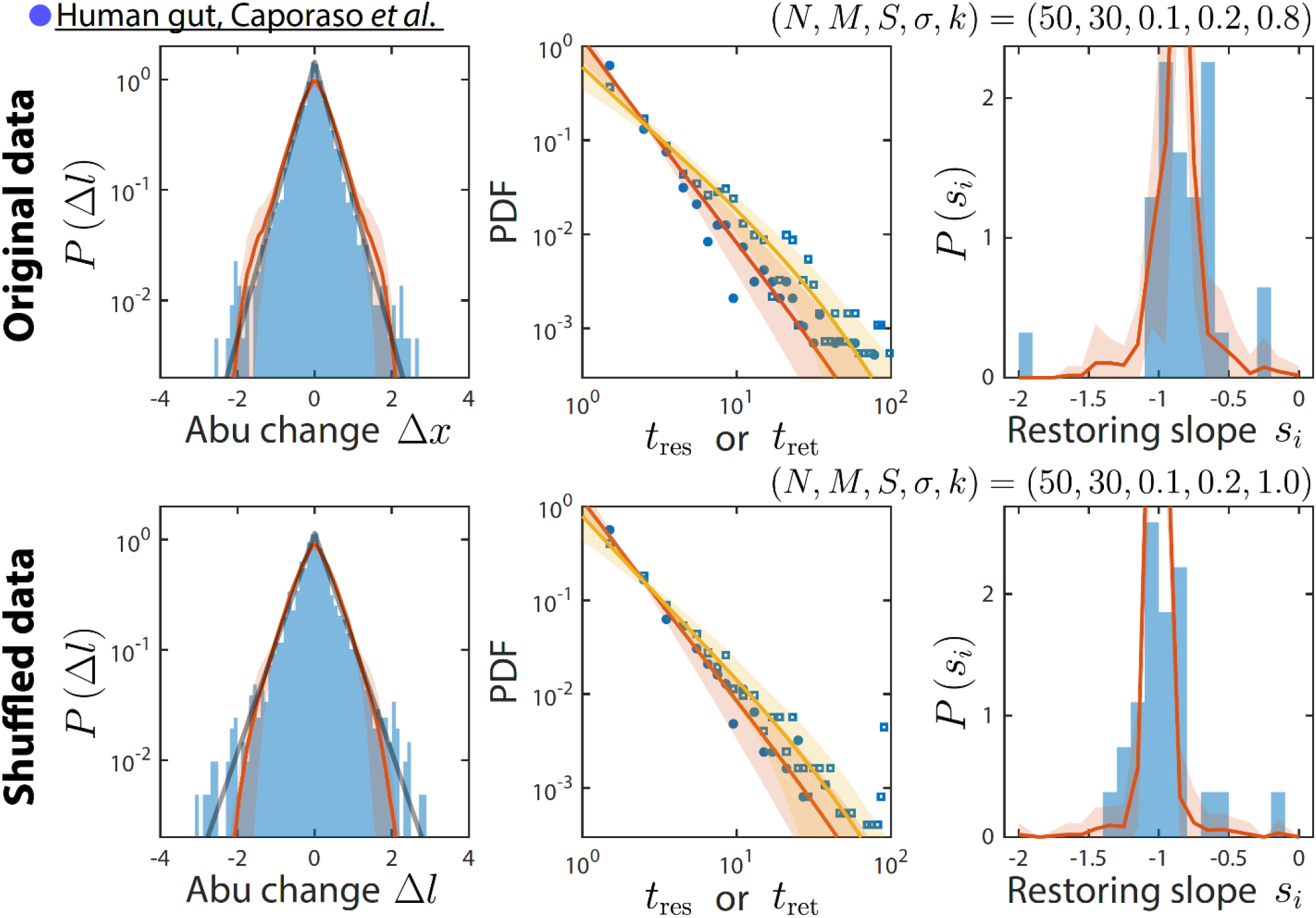
Our mechanistic model is consistent with results obtained by shuffling time labels. Shown are the original data from Ref. ^9^ and the best fit model predictions as in Fig. 2 (top), and the same data but with time labels shuffled (bottom). Only time series statistics that are affected by shuffling time labels are shown. Model predictions are based on the best fit parameters for the shuffled data, which were the same as for the original data except *k* = 1. Shuffling led to more negative restoring slopes, as expected from abolishing correlation between sampling times. The resulting mean restoring slope yielded a best fit value of *k* = 1, which in turn predicted the distributions of residence and return times after shuffling.

**Figure S9:**
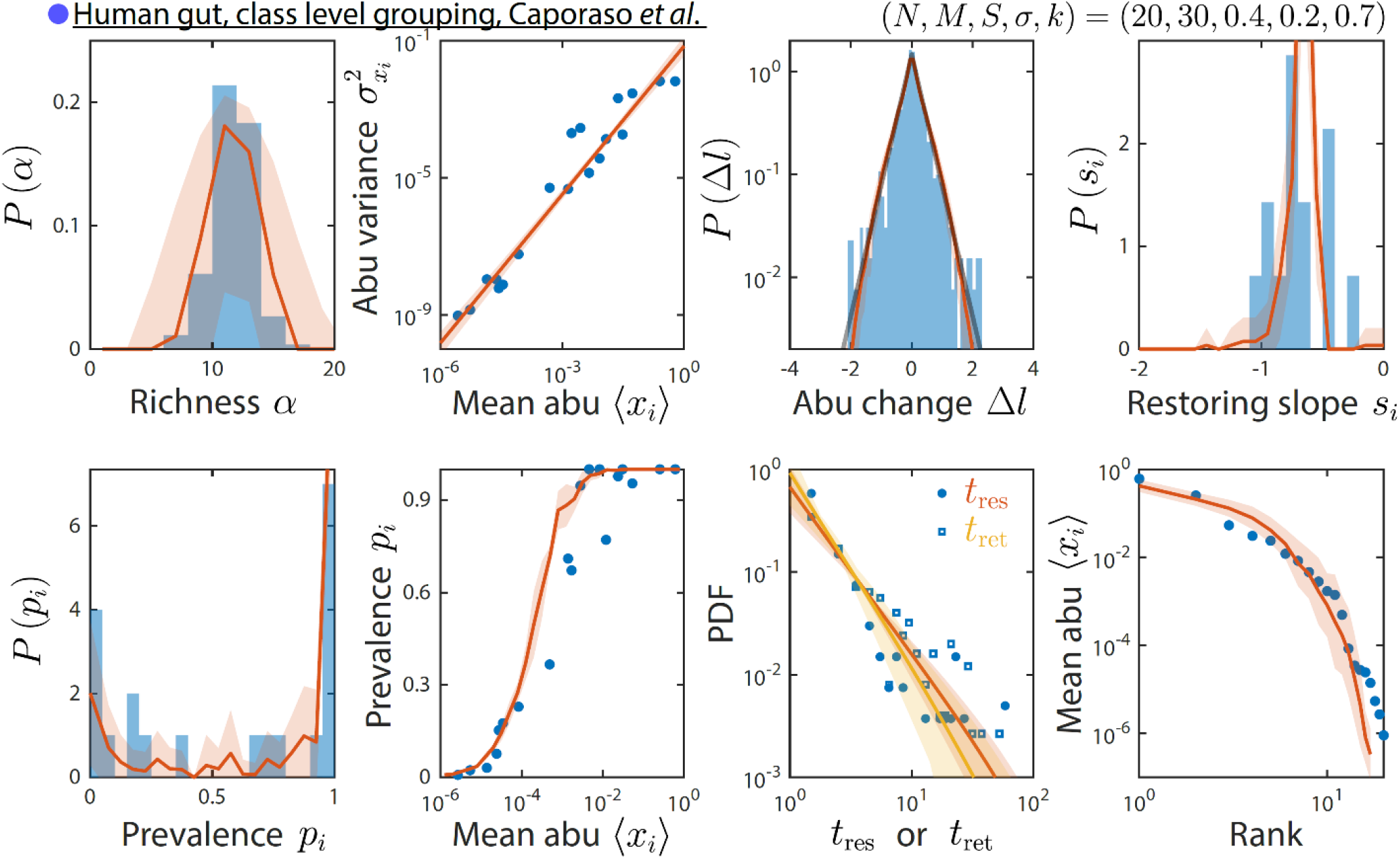
Grouping at a lower-resolution taxonomic level results in similar time series statistics. Shown are data from Ref. ^9^ as in Fig. 2, but grouped and analyzed at the class instead of family level. The best fit model predictions are shown, with best fit parameters (*N, M, S, σ, k*) = (20,30,0.4,0.2,0.6).

**Figure S10:**
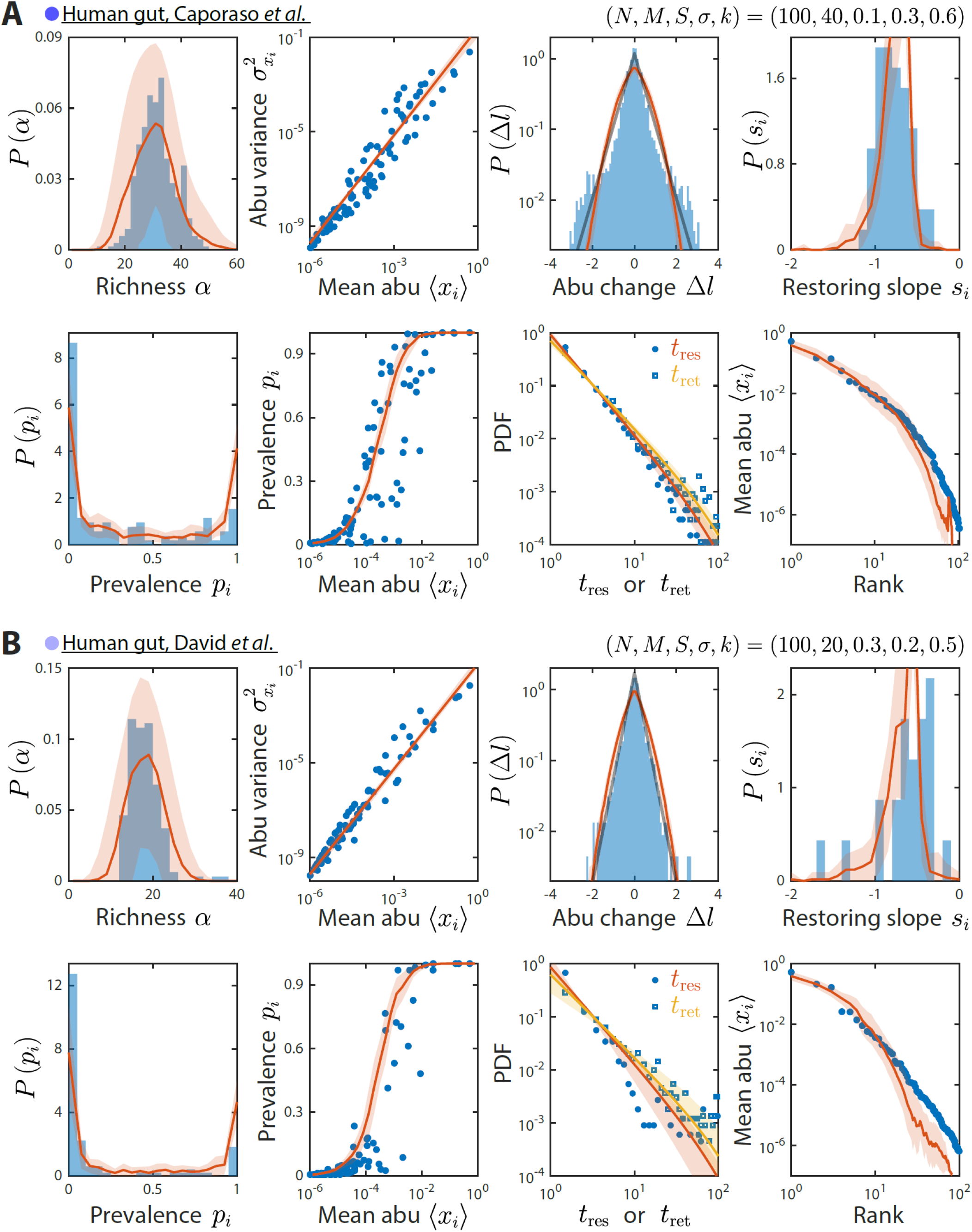
Our model reproduces experimentally observed time series statistics in human gut microbiotas. Shown are the same time series statistics as in Fig. 2 for representative data sets (blue) from Ref. ^9^ (A) and Ref. ^10^ (B) and best fit model predictions (red).

**Figure S11:**
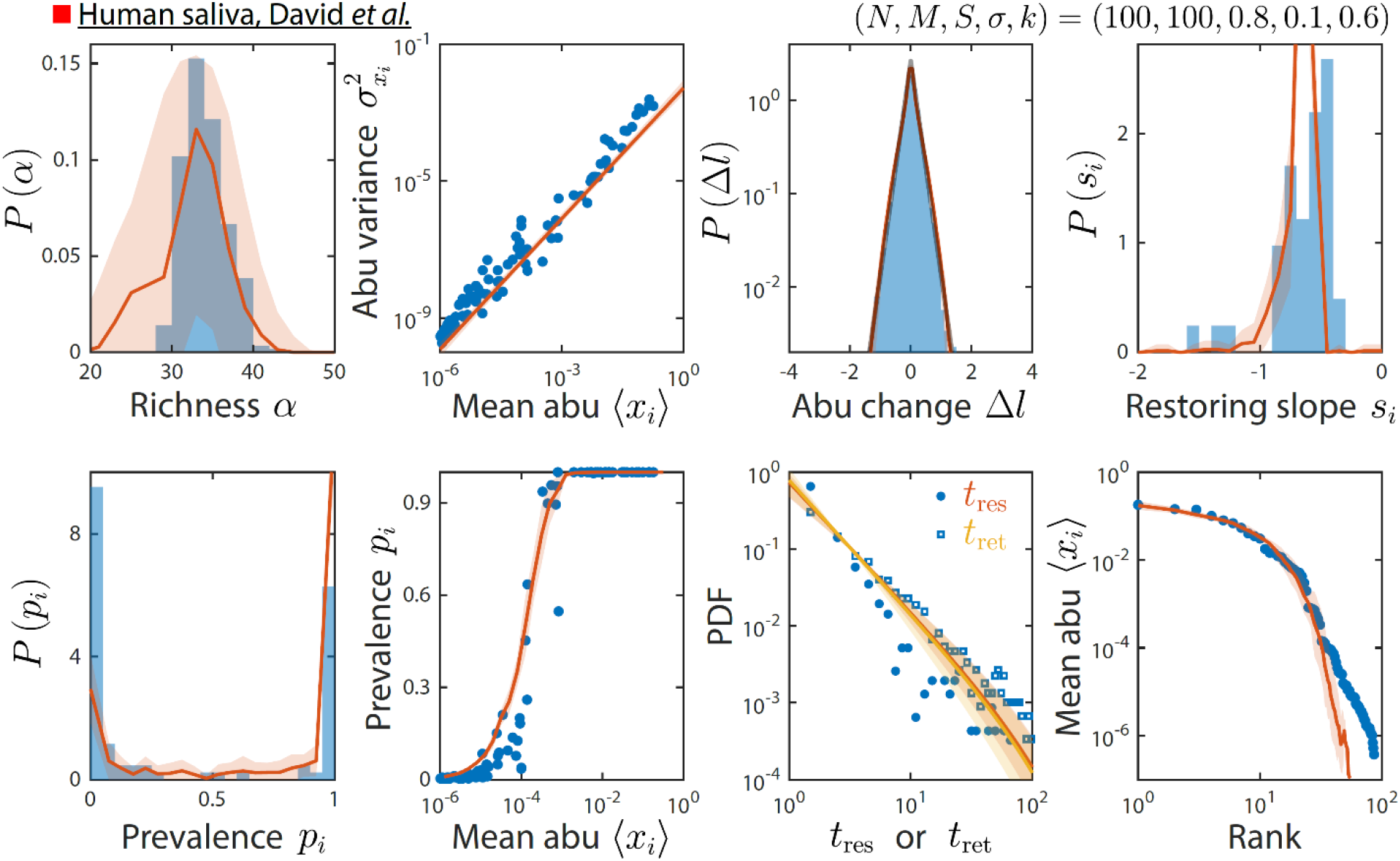
Our model reproduces experimentally observed time series statistics in a human saliva microbiota. Shown are the same time series statistics as in Fig. 2 for an experimental data set (blue) from Ref. ^10^ and best fit model predictions (red).

**Figure S12:**
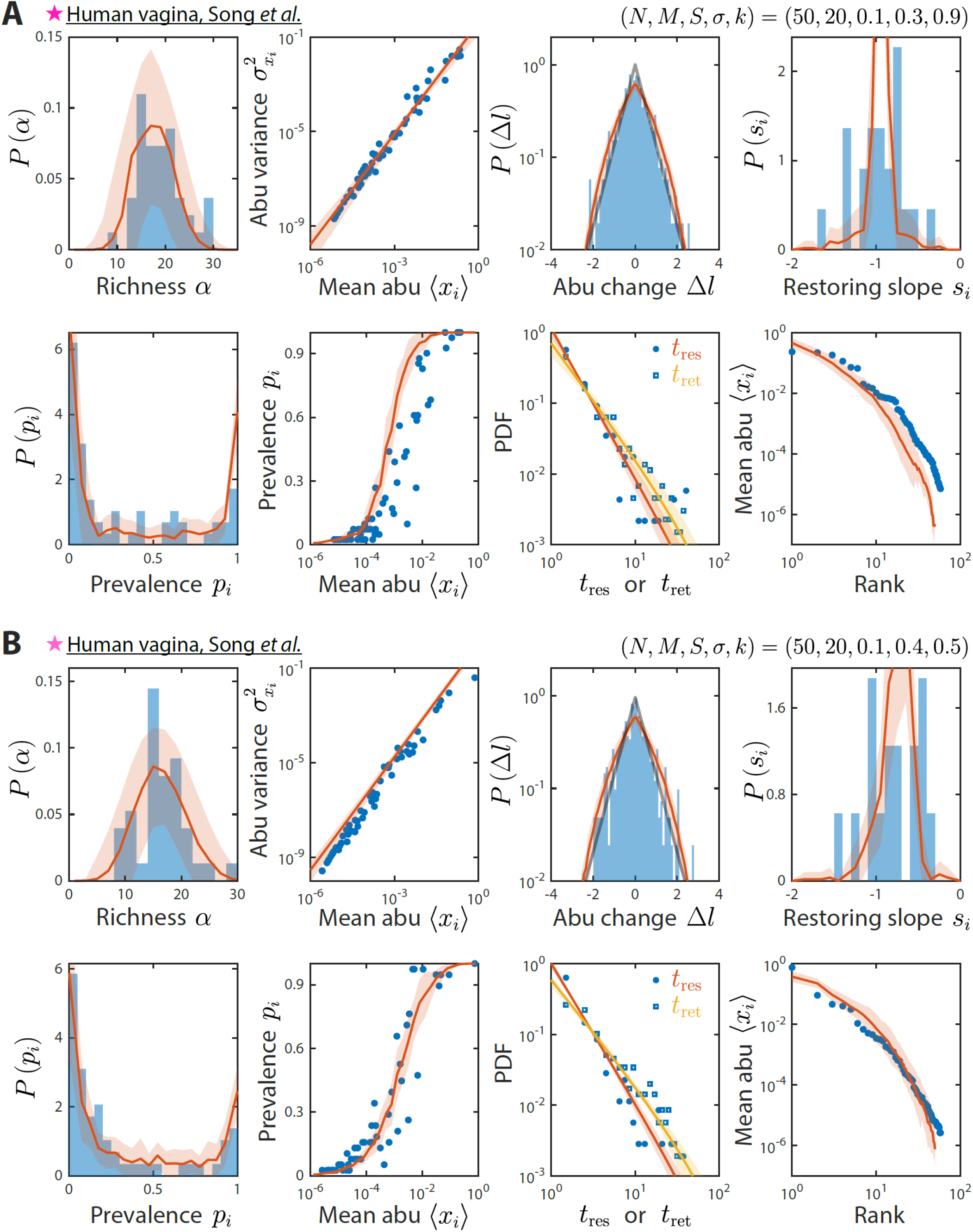
Our model reproduces experimentally observed time series statistics in human vagina microbiotas. Shown are the same time series statistics as in Fig. 2 for representative data sets (blue) of vaginal microbiotas with high diversity (A) and dominated by *Lactobacillus iners* (B) from Ref. ^28^ and best fit model predictions (red).

**Figure S13:**
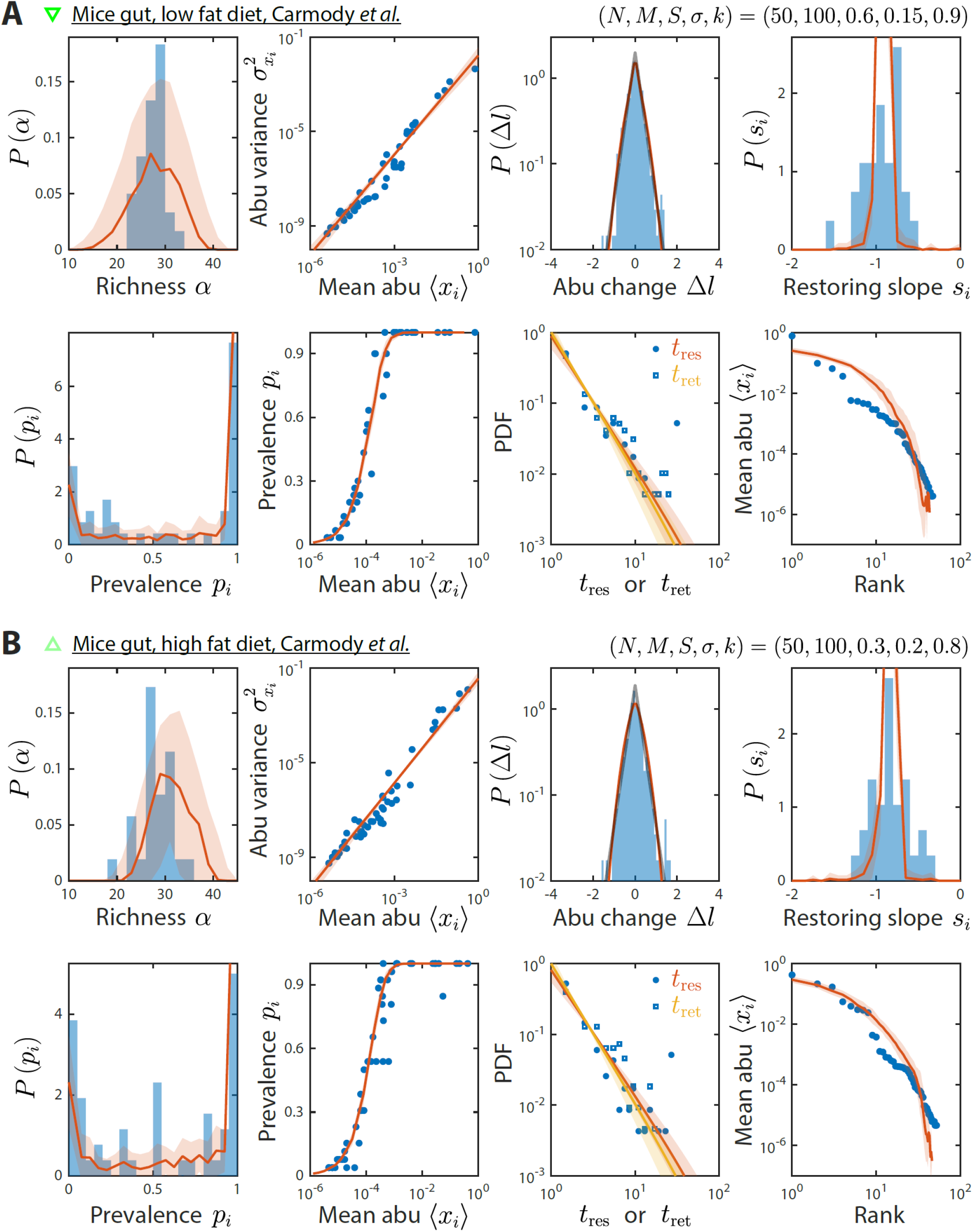
Our model reproduces experimentally observed time series statistics in mice gut microbiotas. Shown are the same time series statistics as in Fig. 2 for representative data sets (blue) of mice fed a low-fat (A) and high-fat diet (B) from Ref. ^27^ and best fit model predictions (red).

**Figure S14:**
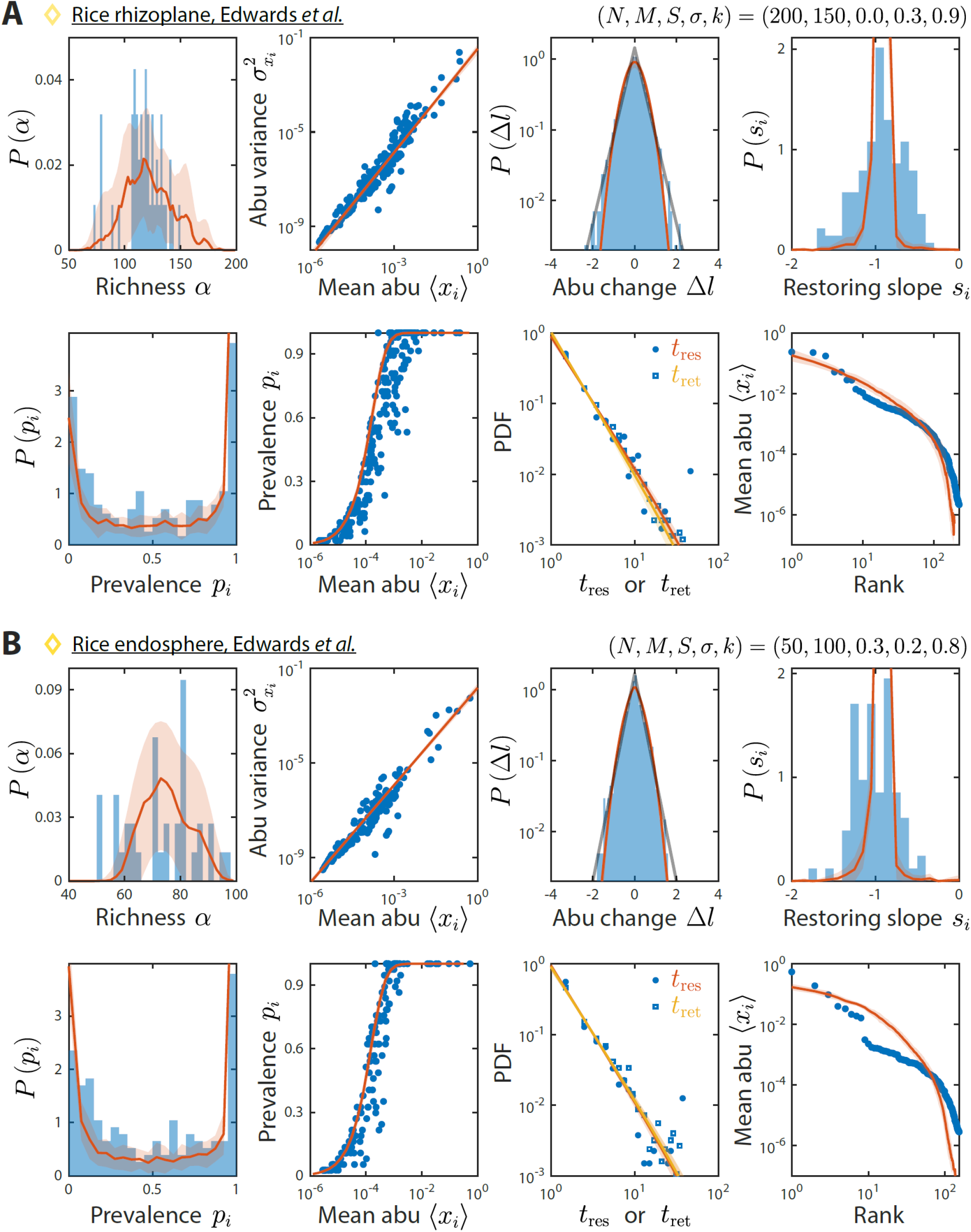
Our model reproduces experimentally observed time series statistics in rice microbiotas. Shown are the same time series statistics as in Fig. 2 for representative data sets (blue) of the rhizoplane (A) and endosphere (B) from Ref. ^29^ and best fit model predictions (red).

**Figure S15:**
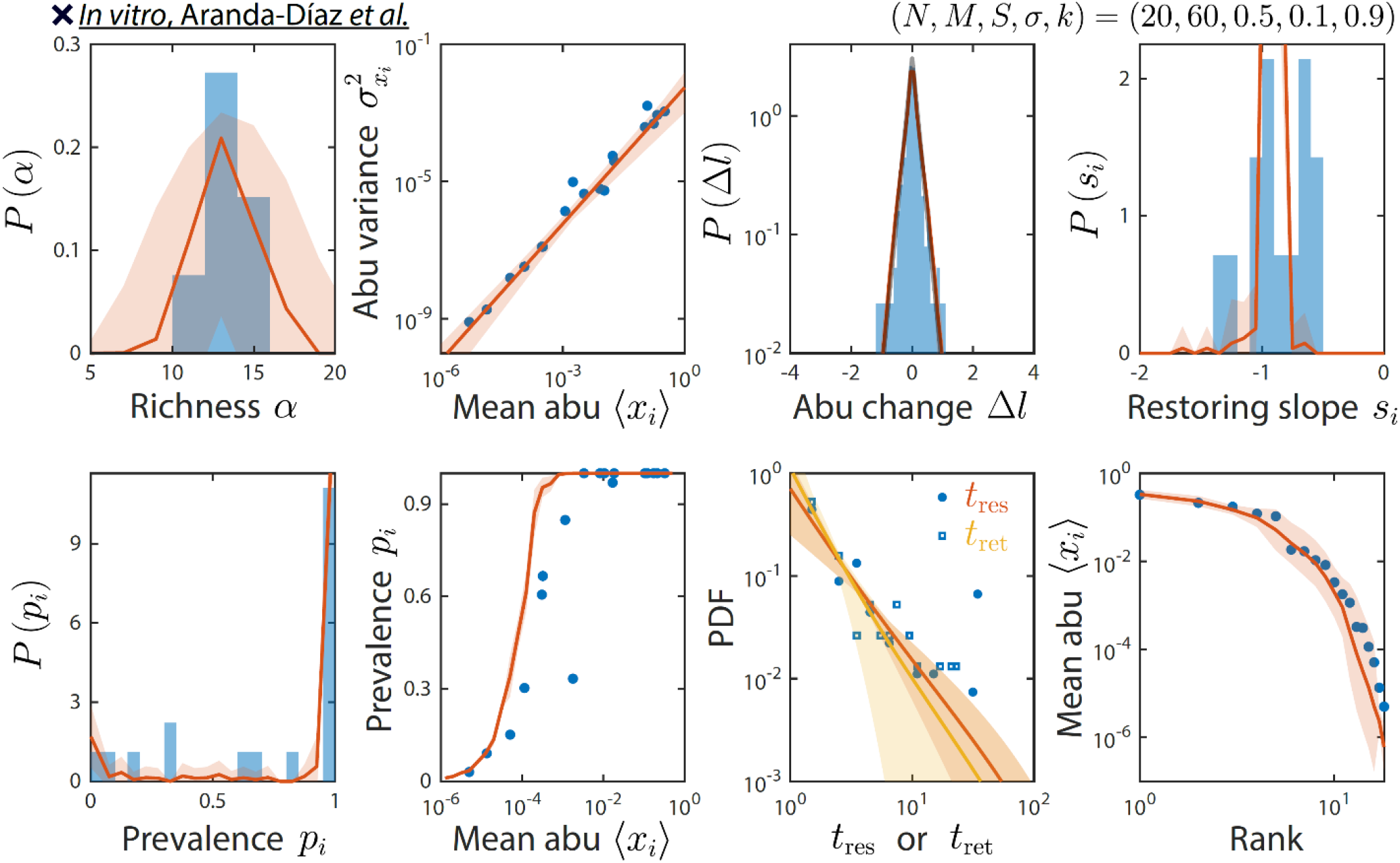
Our model reproduces experimentally observed time series statistics in an *in vitro*-passaged complex community. Shown are the same time series statistics as in Fig. 2 for representative data (blue) from Ref. ^30^ and best fit model predictions (red). Note the lower value of *s* relative to other microbiotas.

## Notes

### Competing Interest Statement

The authors have declared no competing interest.

